# Wilms’ tumor 1 impairs apoptotic clearance of fibroblasts in distal fibrotic lung lesions

**DOI:** 10.1101/2025.05.17.654673

**Authors:** Harshavardhana H. Ediga, Chanukya P. Vemulapalli, Vishwaraj Sontake, Pradeep K. Patel, Hikaru Miyazaki, Dimitry Popov, Martin B Jensen, Anil G. Jegga, Steve K. Huang, Christoph Englert, Andreas Schedl, Nishant Gupta, Francis X. McCormack, Satish K. Madala

**Author notes:** Correspondence: Satish K. Madala, Division of Pulmonary, Critical Care and Sleep Medicine, The University of Cincinnati, CVC 4938, 231 Albert Sabin Way, Cincinnati, OH 45267. **Conflict of interest:** The authors have declared that no conflict of interest exists.

## Abstract

Idiopathic pulmonary fibrosis (IPF) is a fatal fibrotic lung disease characterized by impaired fibroblast clearance and excessive extracellular matrix (ECM) protein production. Wilms’ Tumor 1 (WT1), a transcription factor, is selectively upregulated in IPF fibroblasts. However, the mechanisms by which WT1 contributes to fibroblast accumulation and ECM production remain unknown. Here, we investigated the heterogeneity of WT1-expressing mesenchymal cells using single-nucleus RNA sequencing of distal lung tissues from IPF patients and control donors. WT1 was selectively upregulated in a subset of IPF fibroblasts that co-expressed several pro-survival and ECM genes. The results of both loss-of-function and gain-of-function studies are consistent with a role for WT1 as a positive regulator of pro-survival genes to impair apoptotic clearance and promote ECM production. Fibroblast-specific overexpression of WT1 augmented fibroproliferation, myofibroblast accumulation, and ECM production during bleomycin-induced pulmonary fibrosis in young and aged mice. Together, these findings suggest that targeting WT1 is a promising strategy for attenuating fibroblast expansion and ECM production during fibrogenesis.

## Introduction

Pulmonary fibrosis is the final common pathologic pathway of a variety of acute and chronic lung injuries that are associated with dysregulated healing responses that include myofibroblast accumulation and aberrant lung regeneration (1). Despite its clinical and public health significance, the pathophysiology of pulmonary fibrosis remains incompletely understood. However, it is often associated with fibroproliferation, fibroblast-to-myofibroblast transformation (FMT), and impaired apoptotic clearance of (myo)fibroblasts (2, 3). These processes collectively result in excessive extracellular matrix (ECM) production and scar tissue formation in the lung parenchyma. Idiopathic pulmonary fibrosis (IPF) is perhaps the most severe and enigmatic form of interstitial lung disease, and recent epidemiological studies suggest that the prevalence of the disease is increasing in the U.S. and globally (4, 5). The median survival after diagnosis is poor, typically in the range of 3–5 years (6). The mortality rate of IPF continues to rise each year, particularly among older adults with a history of environmental and occupational exposures (7). It remains one of the most rapidly progressive and fatal forms of interstitial lung disease in the aging population (8). The advent of two FDA-approved therapies for IPF has been a welcome advance for the field, but enthusiasm is tempered by their suppressive rather than remission-inducing effects and their formidable side effect profiles (3, 9, 10). While IPF pathogenesis is extensively studied, the molecular processes underlying impaired fibroblast clearance and ECM production remain largely unknown.

Wilms’ Tumor 1 (WT1), a zinc-finger transcription factor, is a positive regulator of fibroproliferation that promotes excessive collagen and other ECM production in IPF fibroblasts. Several published studies have shown that WT1 is oncogenic in Wilms’ tumors and hematological malignancies (11, 12). WT1 is expressed at high levels in leukemic blast cells, increasing progenitor cell proliferation and survival (13, 14). WT1 is expressed during lung development in most mesothelial cells (and not in myofibroblasts) (3), where it is thought to mediate transformation into fibroblasts and smooth muscle cells in a process called mesothelial-to-mesenchymal transformation (MMT) (3, 15–17). During postnatal and adult stages of lung growth, there is little or no expression of WT1, and MMT does not occur (3, 15, 16, 18). Supporting the critical role for WT1 in lung development and other critical organs such as the heart, homozygous WT1 mutant mice die at E13.5 to E14.5; however, heterozygous WT1 mutant mice with reduced expression of WT1 are viable, fertile, and normal in size (15). WT1 is overexpressed in the distal lung fibroblasts and mesothelial cells in IPF (15, 16, 18). Similarly, in a mouse model of TGFα- induced pulmonary fibrosis, WT1-positive myofibroblasts have been shown to accumulate in thickened subpleural fibrotic lesions due to transformation of fibroblasts to myofibroblasts through MMT (3, 15, 18). WT1 expression is also elevated in fibroblasts from other organs undergoing fibrotic remodeling (19–21). Several recent studies have shown that WT1 induces proliferation, MMT, and ECM production in fibroblasts and that the loss of one allele is sufficient to attenuate both TGFα- and bleomycin-induced pulmonary fibrosis (3). However, the mechanisms by which WT1 contributes to myofibroblast accumulation in pulmonary fibrosis have remained unclear.

In this study, we examined the role of WT1 in (myo)fibroblast survival and ECM production by employing single-nucleus RNA sequencing (snRNA-seq) and fibroblast-specific gain-of-function and loss-of-function mouse models. Our findings identify WT1 as a mediator of fibroblast dysfunction and a potential therapeutic target for fibrosing lung diseases.

## Results

### snRNA-seq of the distal lung cells in normal and IPF lungs

To gain unbiased insight into WT1 expression dynamics and cell type specificity in human lungs, we employed snRNA-seq using the 10X Genomics platform to profile nuclei from 18 IPF and 11 normal donor lung samples, isolated from the distal regions of the right lower lobe (Figure 1A). After randomization of samples during library preparation and sequencing to minimize batch effects, followed by doublet removal, cell, and sample QC, we obtained a total of 100058 single-nucleus transcriptomes from 29 QC-passed samples, including 33057 lung nuclei from control subjects and 67001 lung nuclei from IPF patients using Seurat V4.3.0 in R studio. We performed pre-processing, integration, and clustering using reciprocal PCA (rPCA) integrative analysis, a more conservative approach that preserves distinctions between cells in different biological states. All clusters had acceptable QC metrics, and no cluster was composed of nuclei captured only from individual patients, samples or technical covariates, indicating that data integration was successful and shows disease-related heterogeneity (Figure 1B and Supplemental Figure 1). This captured all major cell types of the lung and identified using canonical markers, COL1A2 for mesenchymal cells, PECAM1 for endothelial cells, EPCAM for epithelial cells, and PTPRC for immune cells (Figure 1C and Supplemental Figure 2). Using unsupervised clustering and canonical markers, we identified 28 unique cell subtype populations, including mesenchymal cells (7), epithelial cells (7), endothelial cells (6), and immune cells (8) (Supplemental Table 1 and Figure 1D). Heatmaps depict the marker gene expression pattern for each major cell type and subject, supporting the validity of our clustering and annotations (Figure 1E). Among the four major cell types, we observed an overall increase in the proportion of mesenchymal, epithelial, and endothelial cells compared to immune cells (22, 23). highlighting the effectiveness of single-nuclei isolation in capturing parenchymal cell types that are traditionally more difficult to extract as single cells from tissues than immune cells (Supplemental Figure 3). Within the mesenchymal cells, we identified 7 subtypes, including alveolar fibroblasts, adventitial fibroblasts, myofibroblasts, pericytes, smooth muscle cells (SMCs), mesothelial cells, and WT1 fibroblasts (Figure 2, A, and B and Supplemental Figures 4 and 6). Among the major mesenchymal cell populations, we observed an increased proportion of WT1 fibroblasts, accompanied by a decrease in the proportion of alveolar fibroblasts, in the distal regions of IPF lungs compared to control subjects (Figure 2, A and B). WT1 expression was elevated in mesenchymal samples of IPF, particularly in WT1 fibroblast subpopulation, a trend also observed in other scRNA-seq datasets (Figure 2, C and D, and Supplementary Figure 5). WT1 fibroblasts highly expressed levels of CTHRC1, POSTN, RUNX1, and several collagen genes, including COL1A1, COL1A2, COL3A1, COL4A5, COL6A3, COL14A1, and COL16A1 in IPF subjects compared to control subjects (Figure 2E). While WT1 fibroblasts are found only in IPF subjects, they are composed of both WT1-positive and WT-negative fibroblasts based on WT1 expression. To further investigate the effects of WT1 expression on fibrosis-associated gene expression, we assessed differentially expressed genes in WT1-positive and negative fibroblasts compared to other mesenchymal cells in IPF subjects and visualized the major gene networks using gene function enrichment analysis. Notably, we observed that genes associated with fibroblast proliferation and apoptosis as the key biological processes enriched in WT1-positive fibroblasts compared to WT1-negative fibroblasts (Figure 2F). Both WT1-positive and negative fibroblasts were enriched for ECM genes compared to other mesenchymal cells in IPF subjects. Collectively, these observations indicate that WT1-positive fibroblasts represent a pathogenic mesenchymal cell population in the distal lung in IPF.

**Figure 1.**
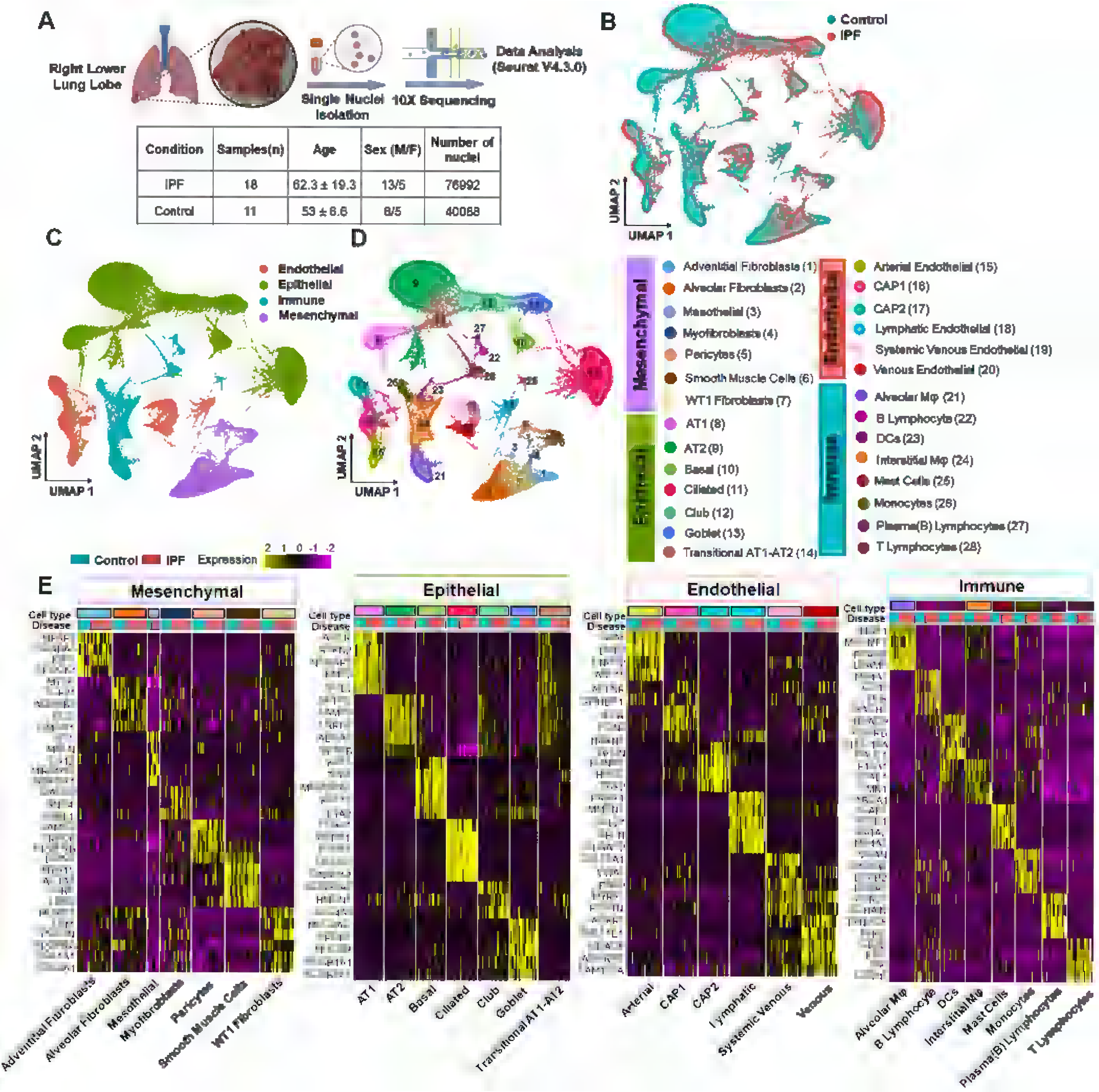
snRNA-seq of lung cells in the distal regions of normal and IPF lungs. **(A)** Schematic of lung single nuclei sample preparation and the workflow for snRNA sequencing and analysis. **(B)** Uniform Manifold Approximation and Projection (UMAP) plot of all cells colored by IPF (n=18) and control donor (n=11) lung samples. Each dot on UMAP represents a nucleus. **(C)** UMAP plot of four major cell lineages colored to denote epithelial, endothelial, mesenchymal, and immune cell types. **(D)** UMAP plot of all 28 unique cell subpopulations identified. **(E)** Heatmaps showing marker gene expression across 28 identified cell types, grouped into four major lineages. Each row represents gene expression across cell types from all samples, while each column shows the average gene expression per subject, grouped by disease condition. Only genes with an adjusted *p*-value < 0.05 (Wilcoxon rank-sum test) are shown. Scaled gene expression values are used for visualization.

**Figure 2.**
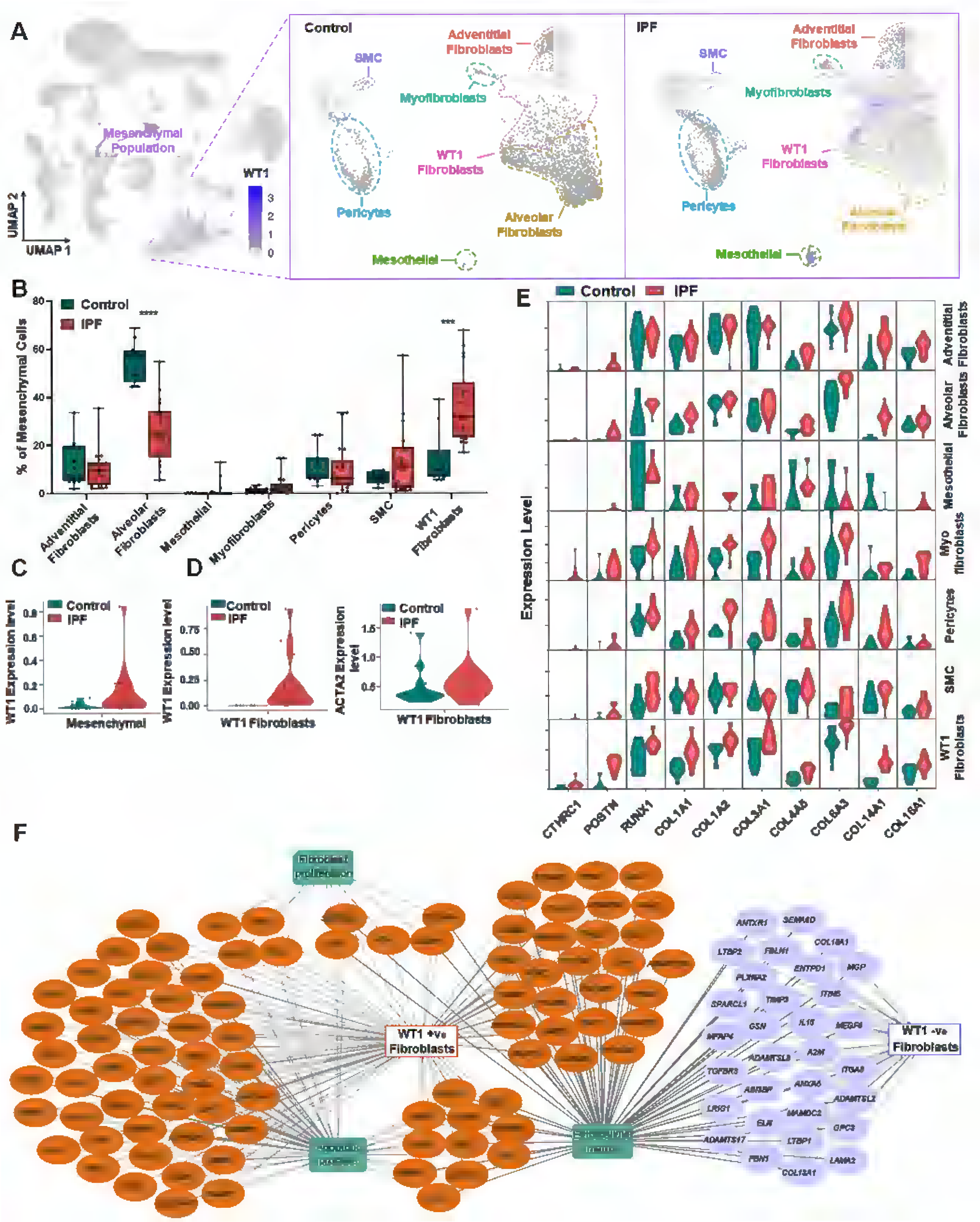
Fibroblast heterogeneity and identification of WT1 fibroblasts in the distal regions of IPF lungs. **(A)** UMAP plots showing normalized WT1 gene expression for all cells and further subdivided by IPF and control donor lung samples with seven mesenchymal cell types. Color intensity reflects expression levels, with darker shades indicating higher expression. **(B)** Boxplots showing the proportion of each cell type within mesenchymal cell types across IPF and control donor lung samples. Each dot represents an individual sample, and the proportions sum to 100% per sample. Multiple t-test was used (\*\*\**p* < 0.001, \*\*\*\**p* < 0.0001). **(C)** Violin plot showing the average WT1 expression in the mesenchymal population, compared between control and IPF samples. Each dot represents an individual sample. The *p*-value (< 0.05) was calculated using the Wilcoxon rank-sum test. **(D)** Violin plot showing the average WT1 expression in the WT1 fibroblasts population, compared between control and IPF samples. Each dot represents an individual sample. The *p*-value (< 0.05) was calculated using the Wilcoxon rank-sum test. **(E)** Violin plots showing the average expression of fibrosis-associated genes (CTHRC1, POSTN, RUNX1, and collagen genes) across all mesenchymal cell subpopulations, comparing samples between control and IPF. **(F)** Differentially Expressed Genes (DEGs) in WT1+ve cells and WT1-ve cells of WT1 fibroblasts population were analyzed using ToppFun and visualized using Cytoscape. Orange- and purple-colored circles represent genes that are upregulated in WT1+ve cells and WT1-ve cells, respectively. The green- colored boxes represent enriched biological processes for the DEGs.

### WT1 is upregulated in distal lung mesenchymal cells in IPF

To validate the accumulation of WT1-expressing mesenchymal cells in the distal regions of IPF lungs, we performed immunostaining for WT1 in the distal lung sections of control donors and IPF subjects. Immunostaining of WT1 supported earlier findings that WT1 is selectively upregulated in mesothelial cells and spindle-shaped mesenchymal cells in fibrotic lesions of pleura and distal lung parenchyma of IPF lungs; however, there was limited or no staining observed in either mesothelial or other lung cells in control subjects (Figure 3A and Supplementary Figure 7).

**Figure 3.**
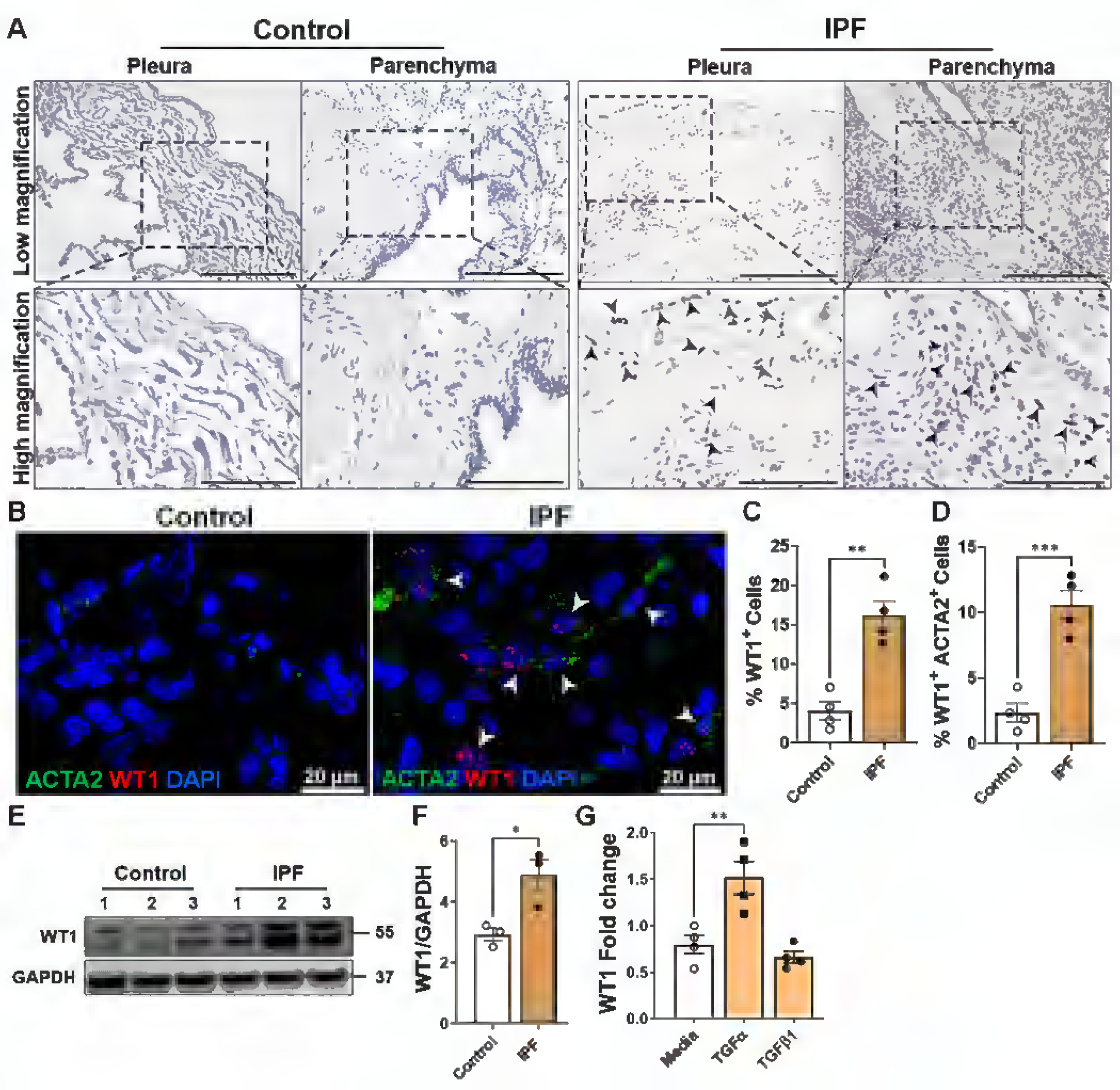
WT1 is upregulated in distal lung mesenchymal cells of IPF. **(A)** Immunostaining was performed with the anti-WT1 antibody on lung sections from control and IPF subjects. Representative images of pleural and parenchymal regions were obtained at low (20X; Scale bar, 200 µm) and high (40X; Scale bar, 100 µm) magnification. Arrowheads highlight WT1 staining on pleural mesothelial cells and spindle-shaped fibroblasts in the distal fibrotic lesions of lung parenchyma. **(B)** Representative fluorescence confocal images for in situ hybridization (RNA-ISH) of WT1 (red) and ACTA2 (green) from control and IPF lungs. White arrowheads illustrate the dual- positive cells. **(C)** Quantification of WT1-positive cells normalized to total lung cells in lung section images from IPF and control lungs. Student’s two-tailed t-test was used (\*\**p* < 0.01; n=4/group). **(D)** Quantification of mesenchymal cells dual positive for WT1 and ACTA2 in the total lung cells in lung section images from IPF and control lungs. Student’s two-tailed t-test was used (\*\*\**p* < 0.001; n=4/group). **(E)** Primary lung-resident fibroblasts isolated from lung cultures of control and IPF lungs were immunoblotted with antibodies against WT1 and GAPDH. **(F)** WT1 protein levels were normalized to GAPDH and were shown as fold change using a bar graph. Student’s two-tailed t-test was used (\**p* < 0.05; n=3/group). **(G)** WT1 transcripts were measured by RT-PCR in normal lung fibroblasts treated with media, TGFα (50 ng/mL) or TGFβ1 (20 ng/mL) for 16 hours (\*\**p* < 0.01; n= 4/group; 1-way ANOVA).

To further validate WT1 upregulation in fibroblasts of IPF lungs, we conducted RNA in situ hybridization for WT1 and ACTA2, a well-established myofibroblast marker, in control and IPF lung tissues. Consistent with snRNA-seq findings, we observed a significant increase in mesenchymal cells co-staining for both WT1 and ACTA2 in distal lung regions of IPF but not in control donor lungs (Figure 3B and Supplemental movies 1-3). Quantitative analysis confirmed a marked increase in WT1-positive cells and cells positive for both WT1 and ACTA2 in IPF, supporting their selective accumulation in distal fibrotic lung lesions of IPF (Figure 3, C and D).

To show that WT1 is upregulated in IPF mesenchymal cells, we measured WT1 protein in fibroblasts isolated from IPF and control lungs (Figure 3, E and F). Indeed, we observed a marked increase in WT1 protein levels in IPF fibroblasts compared to fibroblasts isolated from control lungs (Figure 3, E and F). Given that TGFβ and TGFα are known positive regulators of fibroblast activation in the pathogenesis of pulmonary fibrosis (24, 25), we sought to identify the potential role of these growth factors in the upregulation of WT1 in fibroblasts. We treated normal human fibroblasts with media, TGFβ1, or TGFα for 16 hours in low-serum conditions and observed elevated WT1 transcript levels in fibroblasts treated with TGFα but not TGFβ1 compared to media-treated control fibroblasts (Figure 3G). Taken together, these findings suggest that WT1 is upregulated in IPF mesenchymal cells, and TGFα functions as a positive regulator of WT1 expression in lung mesenchymal cells.

### WT1 is a positive regulator of cell survival and ECM gene expression in IPF fibroblasts

To investigate the effects of WT1 on fibroblast survival, we performed the knockdown of WT1 in IPF fibroblasts isolated from the distal fibrotic lung lesions. As shown in Figure 4A, WT1 knockdown led to a significant decrease in the expression of anti-apoptotic genes, including BCL2-L2, BCL-XL, and BCL3, while concurrently increasing the expression of key pro-apoptotic genes such as BAX and BIM. consistent with a role for WT1 in promoting fibroblast survival by modulating apoptotic pathways. Knockdown of WT1 also resulted in the downregulation of ECM genes, specifically COL1α1 and COL6α3, in fibroblasts treated with WT1-specific siRNA compared to controls treated with non-targeting siRNA (Figure 4B). To validate the impact of WT1 deficiency on pro-apoptotic gene expression, we conducted western blot analysis, which revealed a significant upregulation of the apoptosis-inducing protein FAS in IPF fibroblasts following WT1 knockdown (Figure 4, C and D). Conversely, the expression of the anti-apoptotic protein BCL-XL was markedly reduced in WT1-deficient IPF fibroblasts compared to those treated with control siRNA, further confirming the shift towards a pro-apoptotic state in response to WT1 knockdown (Figure 4, C and D). Further, western blot analysis demonstrated a significant reduction in the levels of ECM proteins, including COL1α1, FN1, ELN, and αSMA, in IPF fibroblasts treated with WT1-specific siRNA compared to control siRNA (Figure 4, E-I). Also, we assessed the effects of WT1 knockdown on collagen secretion and found a significant reduction in secreted collagen proteins in conditioned media from IPF fibroblasts treated with WT1-specific siRNA compared to IPF fibroblasts treated with control siRNA (Supplementary Figure 8). These findings highlight the pivotal role of WT1 in supporting fibroblast survival and ECM production, key processes in the pathogenesis of lung fibrosis. To further elucidate the role of WT1 in the impaired clearance of IPF fibroblasts, we conducted TUNEL assays on IPF fibroblasts treated with either control siRNA or WT1-specific siRNA for 72 hours. WT1 knockdown significantly increased the number of TUNEL-positive fibroblasts under basal conditions and following stimulation with an anti-Fas antibody, compared to cells treated with control siRNA (Figure 4, J and K). These results suggest that WT1 plays a pivotal role in the persistence of IPF fibroblasts by protecting them against apoptotic signals.

**Figure 4.**
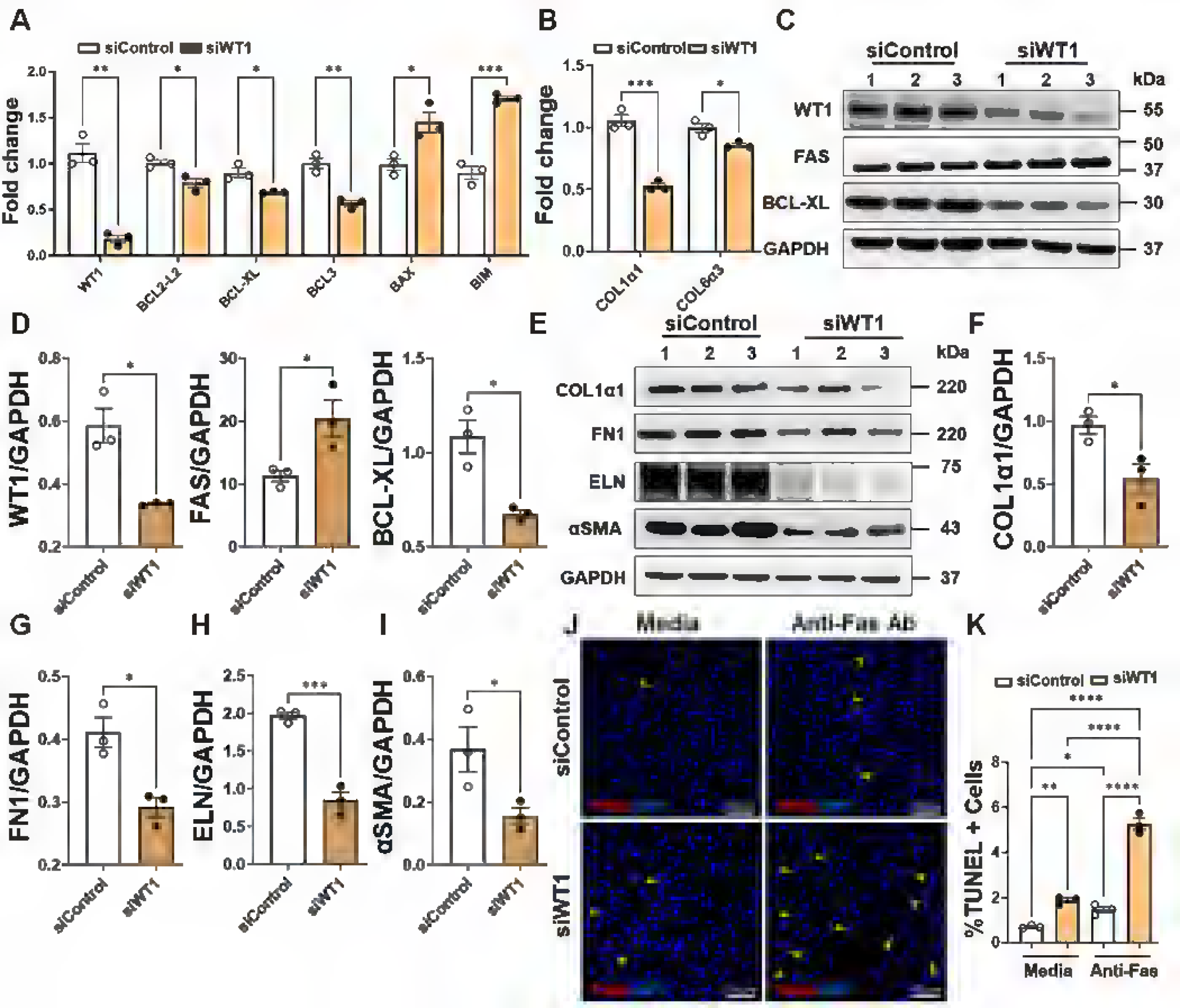
The loss of WT1 attenuates the expression of genes involved in fibroblast survival and ECM production. **(A)** Quantification of both pro-apoptotic (BAX and BIM) and anti-apoptotic (BCL2-L2, BCL-XL, and BCL3) gene transcripts using RT-PCR in IPF fibroblasts treated with either control or WT1 specific siRNA for 72 hr. Multiple unpaired *t*-test were used for comparisons (\**p* < 0.05, ** *p* < 0.01, \*\**p* < 0.001; n= 3/group). **(B)** Quantification of collagen gene transcripts using RT-PCR in IPF fibroblasts treated with either control or WT1-specific siRNA for 72 hr. Multiple unpaired *t*-test was used (\**p* < 0.05, \*\*\**p* < 0.001; n= 3/group). **(C-D)** IPF fibroblasts were treated with either control or WT1-specific siRNA for 72 hr, and cell lysates were immunoblotted with antibodies against WT1, FAS, BCL-XL, and GAPDH. Quantification of WT1, FAS, and BCL-XL protein levels normalized to GAPDH. Student’s two-tailed *t*-test was used (\**p* < 0.05; n=3/group). **(E-I)** IPF fibroblasts were treated with either control or WT1-specific siRNA for 72 hr, and cell lysates were immunoblotted with antibodies against COL1α1, FN1, ELN, αSMA, and GAPDH. Quantification of COL1α1, FN1, ELN, and αSMA protein levels normalized to GAPDH. Student’s two-tailed *t*-test was used (\**p* < 0.05, \*\*\**p* < 0.001; n=3/group). **(J)** IPF fibroblasts were treated with either control or WT1-specific siRNA for 48 hr, followed by anti-Fas treatment for another 24 hr, and cells were stained to quantify total TUNEL positive cells. Representative confocal images were obtained at 20X original magnification. Scale bar: 100 µm. **(K)** The percentage of TUNEL positive (red color) cells in the total DAPI-stained (blue color) cells were quantified using MetaMorph image analysis. One-way ANOVA was used for comparisons (\**p* < 0.05, \*\**p* < 0.01, \*\*\*\**p* < 0.0001; n= 3/group).

### WT1 overexpression promotes fibroblast survival and ECM production in normal fibroblasts

To determine whether WT1 overexpression alone can promote survival and ECM gene expression, we treated normal lung fibroblasts with either a control adenovirus or a WT1- overexpressing adenovirus for 72 hours. As expected, we observed a significant increase in WT1 transcript levels in fibroblasts infected with the WT1-expressing adenovirus compared to the control adenovirus (Supplementary Figure 9). To examine the effects of WT1 overexpression, we selected a panel of pro-survival genes that are differentially expressed in WT1-positive myofibroblasts in our snRNA-seq datasets. We confirmed that overexpression of WT1 induces the expression of pro-survival genes, including HSP90B1, and PIM1 (Figure 5A). Similarly, WT1 overexpression in normal lung fibroblasts resulted in a marked upregulation of anti-apoptotic genes, including BCL2, BCL2-L2, BCL-XL, and BCL3, mirroring the inverse effects observed in WT1 loss-of-function studies in IPF fibroblasts (Figure, 5B and 4A). Also, WT1 overexpression resulted in a significant increase in the expression of several ECM-associated gene transcripts, including COL1A1, COL6A3, and COL16A1 (Figure 5C). We next confirmed the effects of WT1 overexpression on BAD, BAX, FAS, and αSMA protein levels by western blot analysis of normal fibroblasts infected with WT1-overexpressing adenovirus compared to control adenovirus for 72 hours. We observed a significant decrease in the expression of pro-apoptotic proteins BAD, BAX, and FAS and an increase in αSMA levels in WT1- overexpressing fibroblasts (Figure 5, D-I). To determine whether WT1 induces fibroblast survival, we performed TUNEL assays in normal fibroblasts treated with either control adenovirus or WT1-overexpressing adenovirus for 72 hours. WT1 overexpression in normal fibroblasts resulted in a significant decrease in TUNEL-positive cells upon treatment with anti-Fas antibody compared to control (Figure 5, J and K). Collectively, these findings suggest that WT1 functions as a negative regulator of fibroblast apoptosis, consistent with a role for WT1 in fibroblast survival, accumulation, and excess ECM production in the pathogenesis of pulmonary fibrosis.

**Figure 5.**
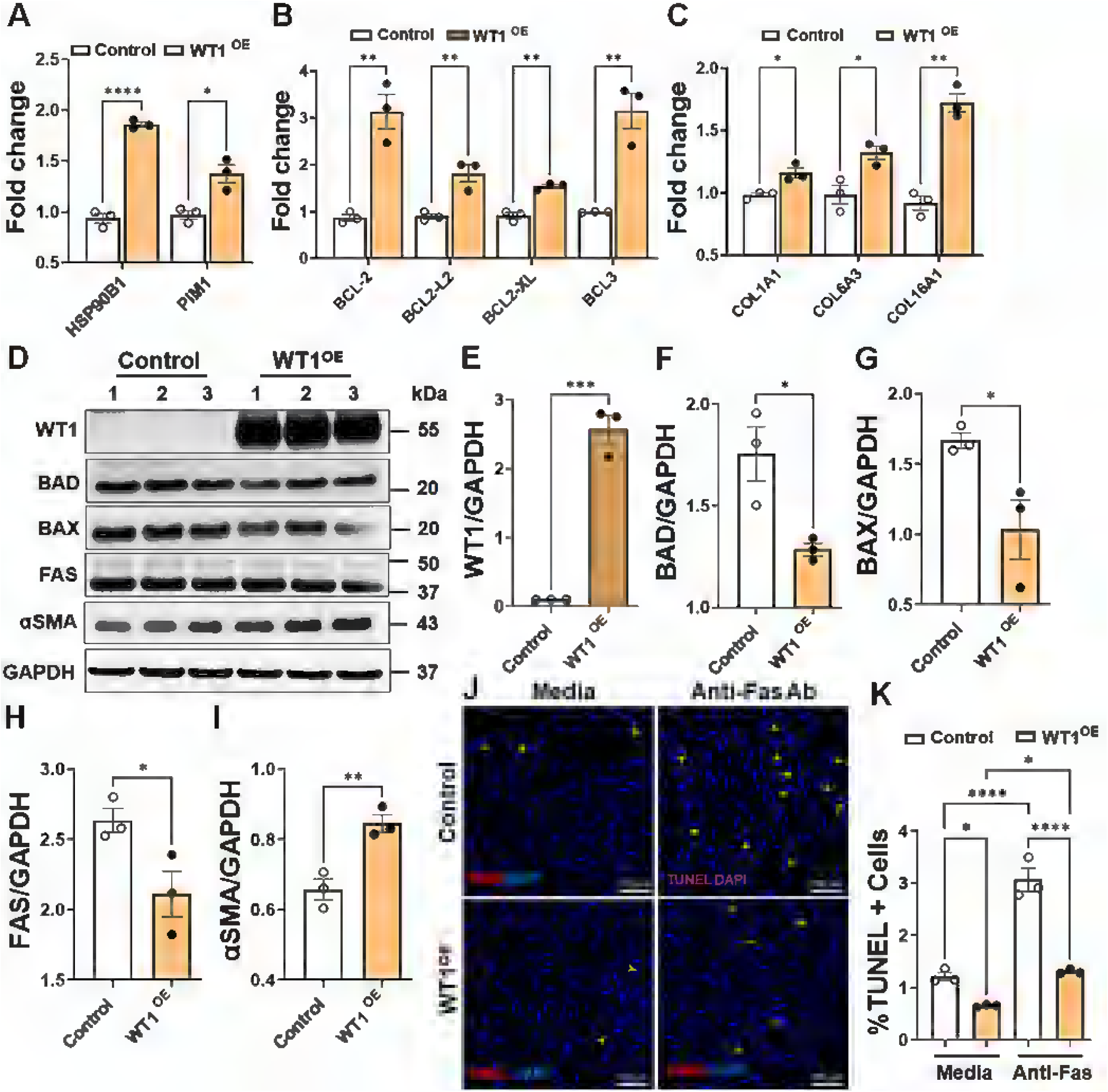
WT1 functions as a positive regulator of lung resident fibroblast survival. **(A)** Quantification of pro-survival gene (HSP90B1, and PIM1) transcripts using RT-PCR in normal lung fibroblasts transduced with either control or WT1 overexpressing adenoviral particles for 72 hr. Multiple unpaired *t*-test were used for comparisons (\**p* < 0.05, \*\**p* < 0.01, \*\*\**p* < 0.001, \*\*\*\**p* < 0.0001; n= 3/group). **(B)** Normal lung fibroblasts were transduced with either control or WT1 overexpressing adenoviral particles for 72 hr and quantified pro-survival gene (BCL2, BCL2-L2, BCL-XL, and BCL3) transcript levels using RT-PCR. Multiple unpaired *t*-test was used (\*\**p* < 0.01; n= 3/group). **(C)** Normal lung fibroblasts were transduced with either control or WT1 overexpressing adenoviral particles for 72 hr and quantified ECM associated gene (COL1α1, COL6α3, COL16α1, and GLI2) transcript levels using RT-PCR. Multiple unpaired *t*-test was used (\**p* < 0.05, \*\**p* < 0.01; n= 3/group). **(D-I)** Normal lung fibroblasts were transduced with either control or WT1 overexpressing adenoviral particles for 72 hr and cell lysates were immunoblotted with antibodies against WT1, BAD, BAX, FAS, αSMA, and GAPDH. Quantification of WT1, BAD, BAX, FAS and αSMA protein levels normalized to GAPDH. Student’s two- tailed *t*-test was used (\**p* < 0.05, \*\*\**p* < 0.001; n=3/group). **(J)** Normal lung fibroblasts were treated with either control or WT1 overexpressing adenoviral particles for 48 hr, followed by anti-Fas treatment for another 24 hr, and fibroblasts stained for TUNEL- positive (red) cells. Representative confocal images at 20X magnification with DAPI- stained nuclei (blue). Scale bar; 100 µm. Yellow arrowheads highlight TUNEL-positive apoptotic cells. **(K)** The quantification of TUNEL-positive fibroblasts using MetaMorph image analysis. One-way ANOVA was used (\**p* < 0.05, \*\*\*\**p* < 0.0001; n= 3/group).

**Figure 6.**
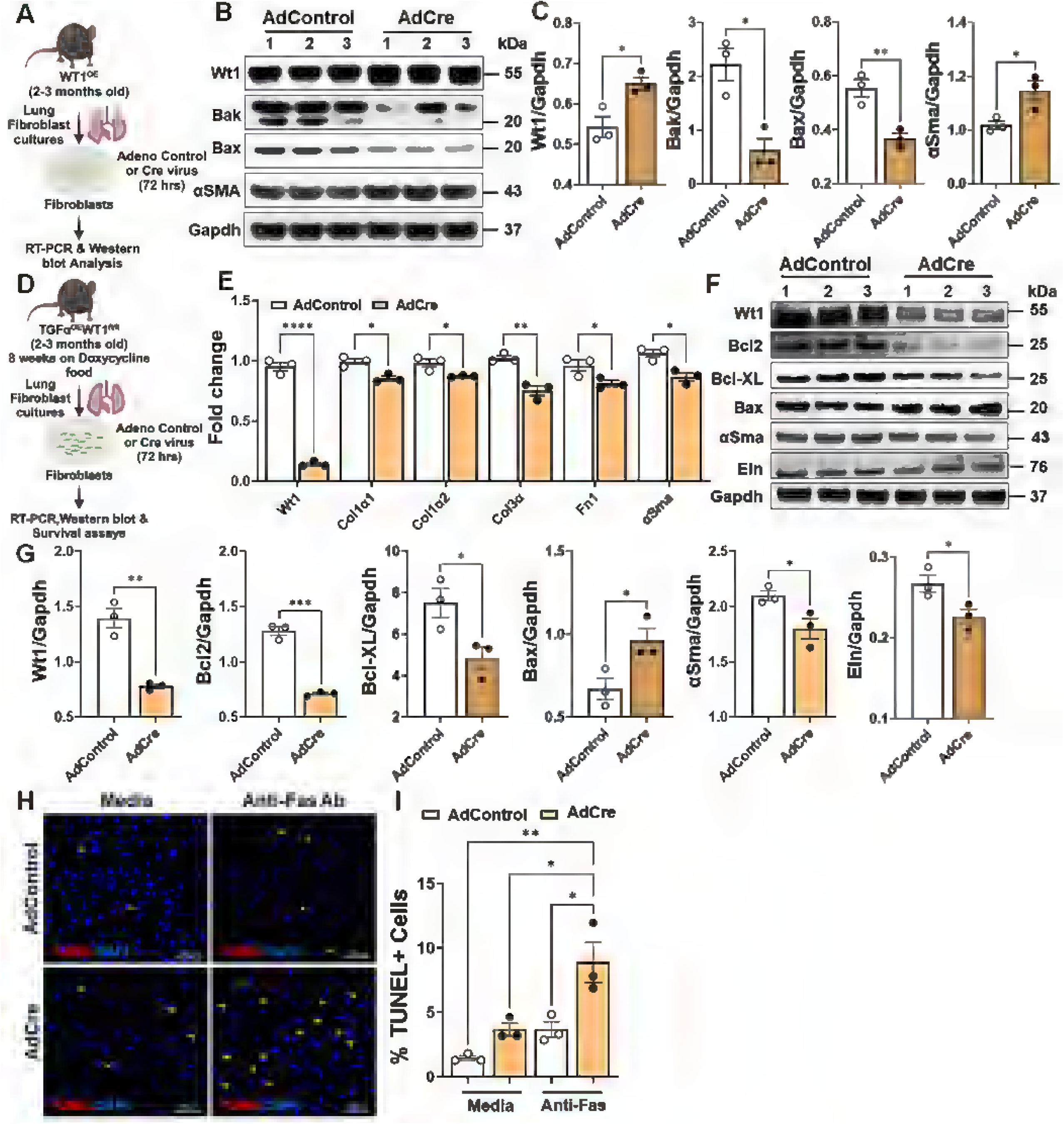
Overexpression of WT1 augments pro-survival gene expression in mouse lung resident fibroblasts. **(A)** Schematic workflow for panels B to C, illustrating the isolation of fibroblasts from lung cultures of WT1^OE^ mice. Fibroblasts were infected with either control adenovirus or Cre-expressing adenovirus for 72 hr to induce WT1 overexpression. **(B)** Fibroblasts were infected with either control adenovirus or Cre- expressing adenovirus for 72 hr, cell lysates were immunoblotted with antibodies against Wt1, Bak, Bax, αSma, and Gapdh. **(C)** Quantification of Wt1, Bak, Bax, and αSma protein levels were normalized to Gapdh. Student’s two-tailed *t*-test was used (\**p* < 0.05, \*\**p* < 0.01; n=3/group). **(D)** Schematic workflow for panels E to I, illustrating the isolation of fibroblasts from lung cultures of TGFα^OE^WT1^fl/fl^ mice on doxycycline food for 8 weeks. Fibroblasts were infected with either control adenovirus or Cre-expressing adenovirus for 72 hr to delete WT1 in fibroblasts isolated from TGFα^OE^WT1^fl/fl^ mice. **(E)** Quantification of WT1 and ECM gene transcripts including Col1α1, Col1α2, Col3α, Fn1, and αSma using RT-PCR in WT1 deficient fibroblasts compared to control fibroblasts. Multiple unpaired t- test was used (\**p* < 0.05, \*\**p* < 0.01; \*\*\*\**p* < 0.0001; n= 3/group). **(F)** Lung–resident fibroblasts isolated from lung cultures of TGFα^OE^WT1^fl/fl^ mice on doxycycline food for 8 weeks were infected with either control or Cre-expressing adenovirus for 72 hr, and cell lysates were immunoblotted with antibodies against Wt1, Bcl2, Bcl-XL, Bax, αSma, Eln, and Gapdh. **(G)** Quantification of Wt1, Bcl2, Bcl-XL, Bax, αSma, and Eln protein levels normalized to Gapdh. Student’s two-tailed *t*-test was used (\**p* < 0.05, \*\**p* < 0.01, \*\*\**p* < 0.001; n=3/group). **(H)** Fibroblasts were infected with either control adenovirus or Cre- expressing adenovirus for 48 hr, followed by treatment with anti-Fas for an additional 24 hr. Cells were then stained to quantify TUNEL-positive apoptotic cells. Representative confocal images were captured at 20X magnification, with a scale bar of 100 µm. Yellow arrowheads point to TUNEL-positive apoptotic cells. **(I)** The quantification of the percent TUNEL-positive cells (red) in the total cells stained with DAPI (blue) was performed utilizing MetaMorph image analysis. One-way ANOVA was used (\**p* < 0.05, \*\**p* < 0.01; n= 3/group).

### WT1 induces survival gene expression in mouse lung fibroblasts

To elucidate the role of WT1 in the survival of lung resident fibroblasts, we generated fibroblasts from the lung cultures of Cre-inducible WT1^OE^ transgenic mice and infected them with control adenovirus or Cre-expressing adenovirus for 72 hours (Figure 6A). As shown in Figure 6B, overexpression of WT1 led to a reduction in the protein levels of the pro-apoptotic markers Bak and Bax, while notably increasing the expression of αSMA compared to control fibroblasts (Figure 6, B and C), suggesting a role for WT1 in promoting fibroblast survival and myofibroblast differentiation.

To examine whether WT1 is required for the maintenance of fibroblast activation, we isolated activated fibroblasts from lung cultures derived from the fibrotic lung lesions of TGFα^OE^ WT1^fl/fl^ mice on Dox for 8 weeks (Figure 6D). We selectively deleted WT1 in activated fibroblasts by treating them with Cre-expressing adenovirus and assessed the changes in survival and ECM gene expression, and apoptosis. As expected, we observed the loss of WT1 in activated fibroblasts by treating them with Cre-expressing adenovirus compared to those in fibroblasts treated with control adenovirus for 72 hours (Figure 6E). The knockdown of WT1 resulted in reduced expression of ECM genes, including Col1α1, Col1α2, Col3α, Fn1, and αSma in activated fibroblasts infected with Cre-expressing adenovirus compared to control adenovirus (Figure 6E). Western blot analysis shows that the knockdown of WT1 resulted in a significant decrease in survival-associated genes including Bcl2 and Bcl-XL, while upregulated the pro-apoptotic Bax protein (Figure 6, F and G). Similarly, WT1 knockdown led to a significant reduction in elastin and αSma protein expression levels compared to control cells (Figure 6, F and G). Additionally, the TUNEL assays demonstrated that WT1 knockdown increased the number of TUNEL-positive fibroblasts in the presence of anti-Fas antibodies, further supporting this finding (Figure 5, H and I). Thus, our findings provide complementary evidence that WT1 is required for the maintenance of the profibrotic functions of activated fibroblasts in fibrotic lung lesions.

### Overexpression of WT1 in fibroblasts augments bleomycin-induced pulmonary fibrosis

To assess the direct effects of WT1 overexpression in fibroblasts, we bred PDGFRα^CreERT^ mice with WT1^OE^ mice to generate fibroblast-specific conditional WT1-overexpression mice (cWT1^OE^) and control mice (PDGFRα^CreERT^). cWT1^OE^ and control (14 week old) mice were treated with two tamoxifen injections per week for 2 weeks. We observed no significant changes in lung architecture or collagen abundance in Masson’s trichrome-stained lung sections from control or cWT1^OE^ mice treated with tamoxifen (Supplementary Figure 10A). and there were no differences in total lung hydroxyproline levels (Supplementary Figure 10B). To assess the upregulation of WT1 and fibrosis-associated genes, we quantified transcript levels of WT1, ECM, and apoptosis-associated genes in total lung RNA from tamoxifen-treated control and cWT1^OE^ mice. As expected, WT1 expression was significantly increased in cWT1^OE^ mice compared to controls. However, no significant differences were observed in the expression of ECM-related genes (Col1α1, Col3α, Col5α, and αSma) or apoptosis-associated genes (Bak1, Bax, Bcl2, and Bcl-XL) between the two groups (Supplementary Figure 10, C-E).

To investigate whether WT1 overexpression augments bleomycin-induced pulmonary fibrosis, control and cWT1^OE^ mice were treated with two tamoxifen injections per week for 7 weeks, and three intratracheal instillations of bleomycin at 3-week intervals for a total of 9 weeks (Figure 7A). We observed a significant increase in WT1 transcript levels in the total lung RNA of cWT1^OE^ mice compared to control mice treated with bleomycin (Figure 7B). To evaluate WT1 upregulation in fibroblasts, we co-immunostained lung sections with antibodies against WT1 and vimentin and observed WT1 upregulation in the fibroblasts of cWT1^OE^ mice compared to control mice treated with bleomycin (Figure 7C). Masson’s trichrome staining revealed a substantial increase in collagen staining in the lung sections of the bleomycin-treated cWT1^OE^ group compared to the bleomycin-treated control mice (Figure 7D). We observed a significant increase in the ratio of the fibrotic area to the total scanned area in the lung sections of cWT1^OE^ mice compared to control mice treated with bleomycin (Figure 7E). Importantly, total hydroxyproline levels were elevated in cWT1^OE^ mice compared to control mice treated with bleomycin (Figure 7F). Consistent with the increase in lung collagen levels, we observed a significant increase in lung resistance in cWT1^OE^ mice compared to control mice treated with bleomycin (Figure 7G). Next, we quantified the changes in ECM-associated gene expression in the total lung transcripts of cWT1^OE^ compared to control mice treated with bleomycin. Consistent with the observed changes in collagen levels, we observed a significant increase in the transcripts of ECM genes, including Col1α1, Col3α, Col5α, Eln, and Fn1, in cWT1^OE^ mice compared to the control mice treated with bleomycin (Figure 7H). Consistent with in vitro findings, we observed a significant decrease in the expression of pro-apoptotic genes, including Bad and Bak1 in cWT1^OE^ mice compared to the control mice treated with bleomycin (Figure 7, I and J). Also, we observed elevated expression of fibroproliferative gene, Plk1 in cWT1^OE^ mice compared to the control mice treated with bleomycin (Figure 7K). To assess the impact of WT1 overexpression on fibroblast activation, we quantified the proliferative changes and apoptosis in lung fibroblasts isolated from the lung cultures of cWT1^OE^ and control groups treated with bleomycin. As assessed by BrdU incorporation, we observed a significant increase in the proliferation of fibroblasts from cWT1^OE^ mice compared to control mice treated with bleomycin (Figure 7L). Similarly, we observed reduced TUNEL-positive cells with anti-Fas treatment in fibroblasts from cWT1^OE^ mice compared to control mice treated with bleomycin (Figure 7, M and N). Taken together, our findings demonstrate the pro-fibrotic effects of WT1 upregulation in promoting fibroblast proliferation, survival, and ECM production during bleomycin-induced pulmonary fibrosis.

**Figure 7.**
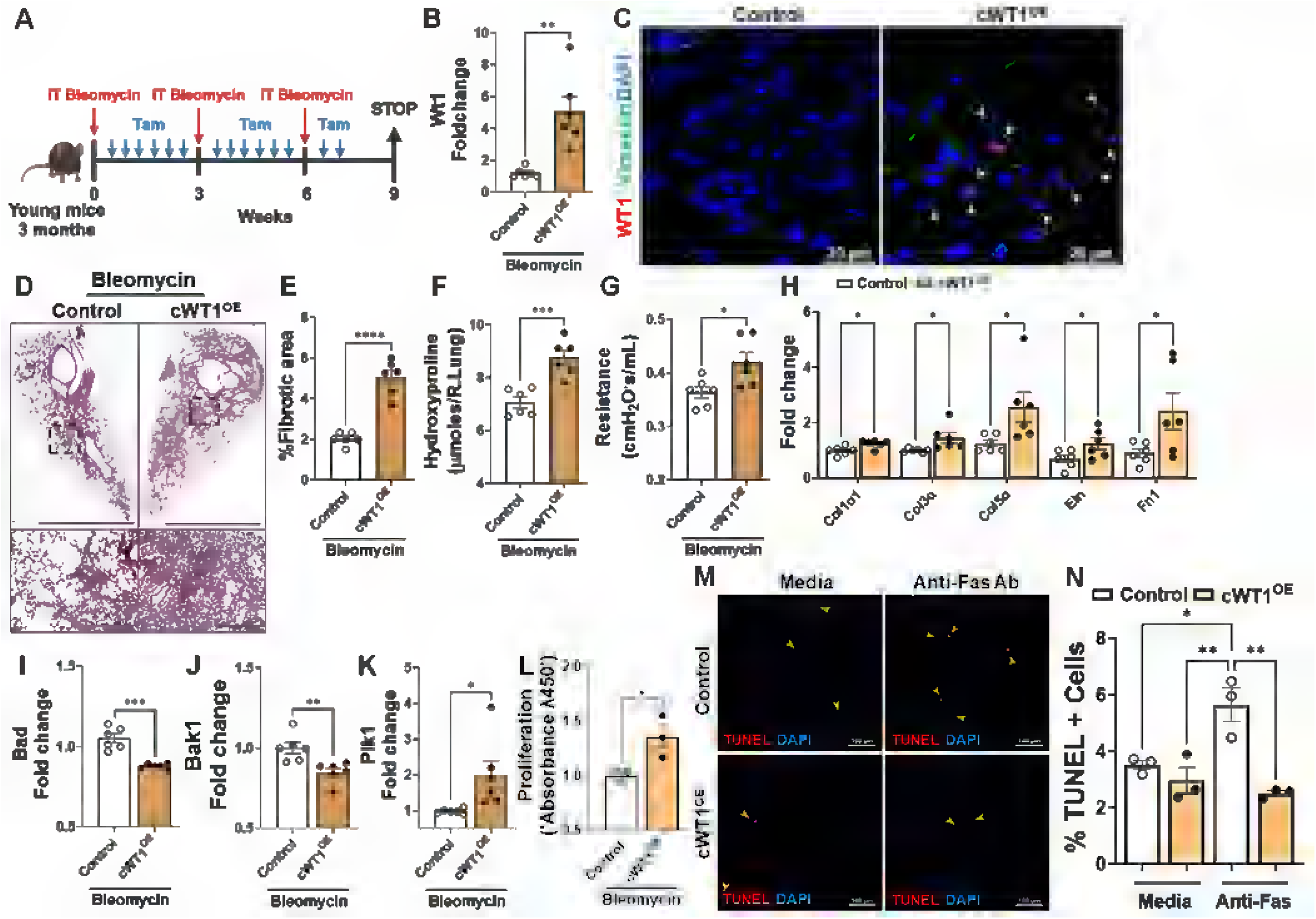
Fibroblast-specific WT1 overexpression augments bleomycin-induced pulmonary fibrosis in mice. **(A)** Schematic representation of the animal experiment involving PDGFRα^CreERT^ (control) and PDGFRα^CreERT^WT1^OE^ (cWT1^OE^) mice. Mice were treated repetitively with bleomycin via intratracheal administration and tamoxifen via intraperitoneal injection as shown in schemata. **(B)** Quantification of WT1 gene transcripts by RT-PCR in the total lung RNA isolated from control and cWT1^OE^ mice. Student’s two- tailed *t*-test was used (\*\**p* < 0.01; n=6/group). **(C)** Representative confocal images of lung sections from control and cWT1^OE^ mice co-immunostained for WT1 (red), vimentin (green) and DAPI (blue). Scale bar: 20 µm. **(D)** Masson’s trichrome-stained lung sections from control and cWT1^OE^ mice. Images were captured at 4X and 20X original magnification with scale bars 1500 µm and 200 µm, respectively. **(E)** Percent fibrotic area was quantified in control and cWT1^OE^ mice using BZ-X image analysis. Student’s two- tailed *t*-test was used (\*\*\**p* < 0.0001; n=6/group). **(F)** Hydroxyproline levels were measured in the right lungs of control and cWT1^OE^ mice. Student’s two-tailed *t*-test was used (\*\*\**p* < 0.001; n=6/group). **(G)** Lung resistance was assessed using Flexivent in both control and cWT1^OE^ mice treated with bleomycin. Student’s two-tailed *t*-test was used (\**p* < 0.01; n=6/group). **(H)** Quantification of ECM gene transcripts (Col1α1, Col3α, Col5α, Eln, and Fn1) from control and cWT1^OE^ mice treated with bleomycin. Multiple unpaired *t*- test was used (* *p* < 0.05; n= 6/group). **(I-J)** Quantification of pro-apoptotic (Bad and Bak) gene transcripts from control and cWT1^OE^ mice treated with bleomycin. Student’s two- tailed *t*-test was used (\*\*\**p* < 0.001; \*\**p* < 0.01; n=6/group). **(K)** Quantification of Plk1 gene transcripts from control and cWT1^OE^ mice treated with bleomycin. Student’s two- tailed *t*-test was used (\**p* < 0.05; n=6/group). **(L)** Proliferation of fibroblasts was quantified using BrdU incorporation assay in fibroblasts isolated from the lung cultures of control and cWT1^OE^ mice treated with bleomycin. Student’s two-tailed *t*-test was used (\**p* < 0.05; n=3/group). **(M)** Fibroblasts isolated from the lung cultures of control and cWT1^OE^ mice treated with bleomycin were cultured and treated with anti-Fas antibody for 24 hr, followed by TUNEL staining (red). Representative confocal images were collected at 20X magnification with DAPI-stained nuclei (blue). Scale bar; 100 µm. **(N)** The percent of TUNEL-positive fibroblasts in total DAPI-positive fibroblasts. One-way ANOVA was used (\**p* < 0.05, \*\**p* < 0.01; n= 3/group). Data are representative of 2 independent experiments with similar findings.

### Overexpression of WT1 in fibroblasts augments pulmonary fibrosis in aged mice

Studying pulmonary fibrosis using aged mice is crucial because aging is a significant risk factor for IPF, and aged mice better replicate the cellular and molecular environment of human IPF, providing a more accurate model for understanding disease mechanisms. Therefore, we overexpressed WT1 selectively in fibroblasts and assessed fibrotic changes in lungs using aged mice with or without bleomycin injury. As shown in Figure 8A, cWT1^OE^ and control mice (15-months-old) were treated with two tamoxifen injections per week, and expression of survival and ECM gene expression were quantified on day 14. Overexpression of WT1 resulted in a significant increase in the expression of WT1 and anti-apoptotic gene expression, including Bcl2, Bcl3, and Bcl-XL (Figure 8B). Further, western blot analysis of total lung lysates supported a significant increase in Col1α1, Bcl2, and Bcl-XL with overexpression of WT1 in cWT1^OE^ mice compared to control mice treated with tamoxifen (Figure 8, C and D). Masson’s trichrome staining exhibited modest or no significant increase in collagen staining in cWT1^OE^ mice compared to control mice following tamoxifen treatment (Supplementary Figure 11, A and B). To assess fibrotic responses with WT1 overexpression during bleomycin-induced injury, cWT1^OE^ and control mice (15-month-old) were treated with bleomycin intratracheally and simultaneously with tamoxifen injections to overexpress WT1 (Figure 8E). Co-immunostaining of lung sections with antibodies against WT1 and vimentin revealed upregulation of WT1 in fibroblasts of cWT1^OE^ mice compared to control mice treated with bleomycin (Figure 8F). We assessed collagen staining in lung sections using Masson’s trichrome staining and observed a significant increase in collagen staining in cWT1^OE^ mice compared to control mice treated with bleomycin (Figure 8, G and H). Notably, we found that the total hydroxyproline levels were elevated in cWT1^OE^ mice compared to control mice treated with bleomycin (Figure 8I). Lung sections immunostained with antibodies against αSMA revealed a significant increase in αSMA staining area in the lungs of cWT1^OE^ mice compared to bleomycin-treated control mice (Figure 8, J and K), consistent with myofibroblast accumulation. Furthermore, Western blot analysis showed WT1 overexpression significantly increased the expression of the anti-apoptotic protein Bcl2, and the ECM protein elastin in cWT1^OE^ mice compared to control mice treated with bleomycin (Supplementary Figure 12, A and B). Thus, our findings indicate that WT1 overexpression in fibroblasts of aged mice contributes to myofibroblast accumulation and collagen deposition during bleomycin-induced pulmonary fibrosis.

**Figure 8.**
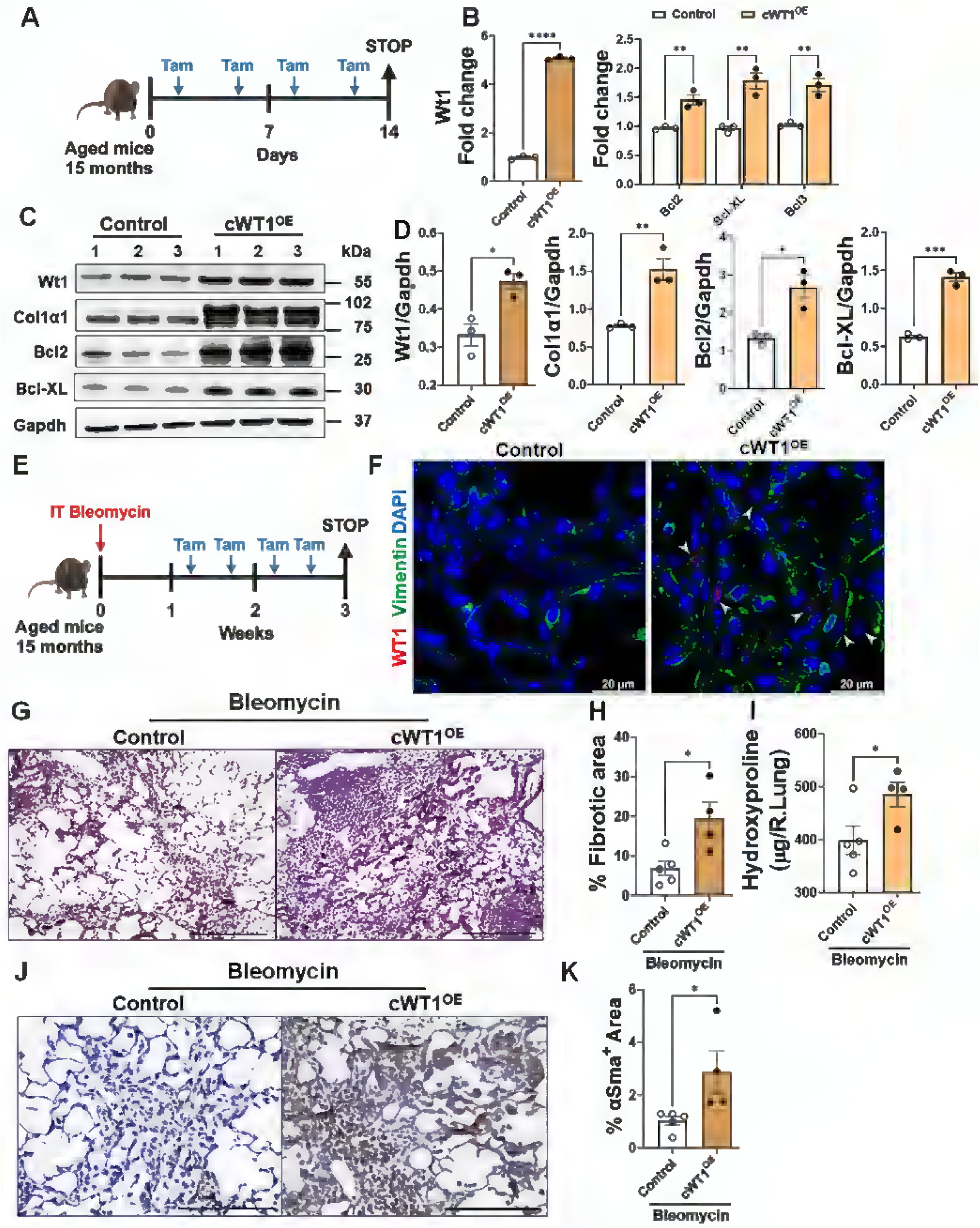
WT1 augments fibroblast survival and bleomycin-induced pulmonary fibrosis in aged mice. **(A)** Schematic presentation of the study using 15 months old PDGFRα^CreERT^ (control) and PDGFRα^CreERT^WT1^OE^ (cWT1^OE^) mice. Mice were treated with two tamoxifen injections per week as shown in schemata. **(B)** Quantification of gene (Wt1, Bcl2, Bcl-XL, and Bcl3) transcripts involved in cell-survival in total lung transcripts of control and cWT1^OE^. Multiple unpaired *t*-test was used (\*\**p* < 0.01, \*\*\*\**p* < 0.0001; n= 3/group). **(C)** The total lung lysates were immunoblotted with antibodies against Wt1, Col1α1, Bax, Bcl2, Bcl-XL and Gapdh from control and cWT1^OE^ mice treated with tamoxifen for two weeks. **(D)** Wt1, Col1α1, Bcl2, and Bcl-XL protein levels were normalized to Gapdh. Student’s two-tailed *t*-test was used (\**p* < 0.05; n=3/group). **(E)** Schematic representation of the animal experiment involving 15 months old PDGFRα^CreERT^ (control) and PDGFRα^CreERT^WT1^OE^ (cWT1^OE^) mice. Mice were treated with a single dose bleomycin and WT1 overexpression in fibroblasts with tamoxifen injections as shown in schemata. **(F)** Representative confocal images of lung sections from 15-month-old control and cWT1^OE^ mice. Arrow heads highlight lung fibroblasts that co-stained for WT1 (red), vimentin (green), and DAPI (blue) in lung sections of control and cWT1^OE^ mice treated with bleomycin. Scale bar, 20 µm. **(G)** Masson’s trichrome stained lung sections from control and cWT1^OE^ mice treated with bleomycin. Images were captured at 20X original magnification. Scale bar, 200 µm. **(H)** Percent fibrotic area was quantified in control and cWT1^OE^ mice using BZ-X image analysis. Student’s two-tailed *t*- test was used (\**p* < 0.05; n=4-5/group). **(I)** Hydroxyproline levels were measured in the right lungs from control and cWT1^OE^ mice treated with bleomycin. Student’s two-tailed *t*- test was used (\**p* < 0.05; n=4-5/group). **(J)** Representative images of αSMA-stained lung sections of control and cWT1^OE^ mice treated with bleomycin. Images were captured at 40X original magnification. Scale bar, 100 µm. **(K)** Quantification of αSma-positive area in whole lung sections. Student’s two-tailed *t*-test was used (\**p* < 0.05; n=4-5/group).

## Discussion

In this study, we used snRNA-seq analyses to characterize matrix-embedded mesenchymal cell populations in the distal areas of IPF lungs. We identified nine mesenchymal subpopulations, including WT1-expressing fibroblasts and myofibroblasts. Although it has been demonstrated that WT1 overexpression promotes fibroproliferation and ECM production, whether WT1 also impairs apoptotic clearance of lung fibroblasts resulting in progressive accumulation of myofibroblasts has not been established. Here we demonstrate that WT1, typically restricted to mesothelial cells during lung development, is aberrantly upregulated along with anti-apoptotic genes in the distal lung fibroblasts in IPF. We conclude that the expression of WT1 in fibroblasts marks a pathogenic shift, contributing to the impaired clearance of fibroblasts and the progression of fibrotic remodeling in the lung.

The observed upregulation of WT1 in IPF fibroblasts suggests a unique and previously unrecognized role for this developmental transcription factor in lung fibrosis. Our in vitro and in vivo findings are consistent with this notion, as immunohistochemical analyses revealed that WT1 is predominantly localized in the nuclei of spindle-shaped cells within fibrotic lesions of the distal lung, and WT1 upregulation was associated with elevated fibroproliferation, ECM production, and resistance to apoptosis-all key processes in the development and maintenance of fibrotic lung lesions. The functional role of WT1 in fibroblast survival and ECM production was further supported by our loss- and gain-of-function studies, as WT1 knockdown in IPF fibroblasts resulted in a marked reduction in ECM-related gene expression and increased sensitivity to apoptosis. In vivo, overexpression of WT1 in fibroblasts exacerbated fibrosis in bleomycin-induced pulmonary fibrosis models, evidenced by increased collagen deposition, myofibroblast accumulation, and impaired lung function. Further, the deletion of WT1 in activated fibroblasts isolated from fibrotic lesions attenuated their proliferation and survival. These complementary in vitro and in vivo experiments establish WT1 as a pivotal regulator of fibroblast survival and fibrosis severity. WT1-expressing fibroblasts that accumulate in the distal lung regions express high levels of pro-survival and anti-apoptotic BCL-2 family genes. The high expression of WT1 in distal fibrotic lesions is consistent with the notion that WT1 may directly provide a cell survival advantage in addition to proliferative and matrix-producing functions. This hypothesis is also supported by the fact that WT1 is required to overcome apoptosis and potentiate proliferation in myeloid leukemia cells (26). Consistent with our data demonstrating that WT1 transcriptionally upregulates the BCL-2, a study by Mayo et al suggests that WT1 positively stimulates the BCL-2 promoter through direct interaction (27). The anti-apoptotic function of WT1 is not limited to fibroblasts in pulmonary fibrosis, as high WT1 expression in osteosarcoma has been shown to promote Bcl2 levels and resistance to apoptosis (28). Similarly, a recent study suggests that WT1 may potentiate oncogenesis and tumor progression by transcriptionally upregulating Bcl2, and that this mechanism may contribute to the heightened resistance to chemotherapy-induced apoptosis in diffuse anaplasia Wilms’ tumors (27).

Our study has several noteworthy limitations. The samples used in our analyses were collected exclusively from the lung periphery, which restricts our ability to investigate the involvement of mesenchymal cell types from other regions of the lung in the pathogenesis of pulmonary fibrosis. However, the use of stored tissue enabled us to process multiple samples simultaneously, minimizing batch effects. While we observed an accumulation of WT1-expressing fibroblasts in the distal regions of IPF lungs, our current methodology does not allow us to determine the extent of crosstalk between WT1- positive cells and other fibroblast subpopulations in these regions, nor their distinct contributions to IPF pathogenesis. Supporting our observations, Habermann et al. identified HAS1-positive, WT1-expressing fibroblasts localized to subpleural regions, characterized by elevated expression of fibrosis-related collagen genes and activation of cellular stress and Th2 cytokine pathways (23).

In this study, we identified a novel role for WT1 in promoting mesenchymal cell survival by regulating the expression of apoptosis-associated genes. This enhanced survival was accompanied by increased collagen deposition and accumulation of myofibroblasts in a bleomycin-induced model of pulmonary fibrosis in both young and aged mice. Notably, in the absence of injury, WT1 overexpression had minimal or no effect on the induction of survival or other profibrotic genes in the lungs of young mice but was sufficient to induce mild fibrosis in aged lungs. Given that IPF is a predominantly age-associated fibrotic lung disease, these findings suggest that age-related epigenetic alterations may have amplified the effects of WT1 in the aged lung. Indeed, previous studies have shown that aging is associated with altered chromatin accessibility, including increased histone acetylation, which could enhance the expression of fibrotic genes and contribute to persistent fibrosis (29). Together, these findings underscore the complex regulatory role of WT1 in fibroblast activation and highlight the need for future studies to dissect the mechanisms governing WT1-driven gene expression in aging tissues.

A study by Karki et al. suggests that WT1 loss in pleural mesothelial cells (PMCs) promotes their migration into the lung parenchyma and mesothelial-to-mesenchymal transformation (MMT). However, more recent studies have demonstrated that WT1 is upregulated in both mesothelial cells and (myo)fibroblasts in IPF, as well as in a mouse model of TGFα-induced pulmonary fibrosis (18, 30). Consistent with these findings, our snRNA-seq and immunostaining analyses reveal increased WT1-positive mesothelial cells and (myo)fibroblasts in IPF. Both in vitro and in vivo studies using loss function and gain of function models demonstrate that WT1 functions as a positive regulator of ECM gene expression, fibroproliferation, myofibroblast transformation, and pulmonary fibrosis (3). While the findings by Karki et al., provide important insights into mesothelial contributions to fibrosis, the broader body of evidence underscores a more complex role for WT1 in fibroblast activation and ECM remodeling. This refined understanding highlights WT1 as a potential therapeutic target in pulmonary fibrosis. While transcription factors have traditionally been challenging to target therapeutically, novel strategies using small-molecule inhibitors and RNA-based approaches are emerging. In particular, several transcription factor-targeting therapies, including inhibitors of AP-1, JAK-STAT, NF-κB, and MYC, are currently under investigation, highlighting the potential feasibility of this approach (31–34). The selective upregulation of WT1 in fibrotic lesions, as opposed to control lung tissue, suggests that WT1 could serve as a promising therapeutic target with limited off-target effects in non-fibrotic tissues. This specificity contrasts with the widespread expression of pro-survival genes such as BCL-2 across multiple lung cell types, which could limit the therapeutic efficacy of BCL2 inhibitors like BH3 mimetics (e.g., ABT-262) due to their off-target effects (35, 36). Our findings thus provide a critical foundation for the development of targeted therapies that leverage the unique expression patterns of WT1 in IPF (3, 18). Indeed, WT1 has emerged as a potential therapeutic target in oncology because of its key role in cell growth and differentiation and limited expression in normal tissues (37, 38). WT1-specific immunotherapies, such as peptide vaccines, dendritic cell vaccines, and adoptive T-cell therapies, are being designed to enhance the immune recognition and destruction of WT1-expressing cancer cells (37, 39). Small-molecule inhibitors targeting WT1 directly or its downstream signaling pathways are also under investigation (9, 40). For example, WT1-specific antisense oligonucleotides have shown potential in preclinical models by reducing WT1 expression and impairing cancer cell growth (26). These approaches are also potentially promising for targeting fibroblasts in pulmonary fibrosis, though more research is needed to validate these strategies. Unlike broad TGF-β pathway inhibitors that affect multiple cell types and immune homeostasis, targeting WT1 may provide a more specific means to disrupt fibroblast activation and survival while minimizing off-target effects. Given its aberrant expression in pulmonary fibrosis, targeted delivery strategies, such as nanoparticle-based siRNA approaches, may help mitigate toxicity concerns (41, 42). Therefore, future studies are warranted to refine WT1-targeting approaches and assess their therapeutic efficacy in IPF and other fibrotic diseases.

In conclusion, our findings demonstrate that WT1 plays a critical role in the pathogenic activation of fibroblasts in IPF. By promoting fibroblast proliferation, survival, and ECM production, WT1 contributes to the fibrotic remodeling that characterizes IPF. Targeting WT1 or its upstream regulators may offer a new therapeutic strategy for mitigating fibrosis and improving outcomes for patients with IPF. Further studies are warranted to explore the therapeutic potential of modulating WT1 expression or activity and to better understand the broader implications of WT1 reactivation in other fibrotic diseases.

## Methods

### Sex as a biological variable

Our study included both male and female human samples, as well as male and female mice. Our findings are similar for both sexes, indicating that the reported results are not sex-specific.

### Human samples

Human lung tissue samples, including those from patients with idiopathic pulmonary fibrosis (IPF) and healthy controls, were obtained through the Translational Pulmonary Science Center (TPSC) at the University of Cincinnati Medical Center. All sample collection and usage procedures were conducted in compliance with ethical guidelines and approved by the University of Cincinnati Institutional Review Board (IRB #2013-8157), ensuring the protection of human subjects and adherence to confidentiality protocols. Lung tissue from individuals without documented lung disease was used as control donor lungs. To safeguard patient anonymity, all samples and associated data were de-identified before being provided to the research team.

### Mouse strains

The generation of ROSA26:WT1-KTS knock-in mice and WT1^fl/fl^ mouse lines have been previously described (43, 44). To generate fibroblasts-specific WT1 overexpression mice (PDGFRα^CreERT^WT1^OE^; cWT1^OE^), WT1 knock-in mice were crossed with PDGFRα^CreERT^ mice (Stock no. 018280, The Jackson Laboratory). WT1 overexpression was induced in adult cWT1^OE^ mice through Cre activation, achieved by administering tamoxifen twice weekly for two or four weeks, starting at 12-14 weeks (young mice) or 15 months (old mice) of age. The generation of doxycycline-inducible and Clara-cell-specific TGFα overexpression mice (TGFα^OE^ mice) has also been previously described (45). We bred TGFα^OE^ mice with WT1^fl/fl^ mice to generate TGFα^OE^WT1^fl/fl^ mice. To induce TGFα overexpression, mice were fed a diet containing doxycycline (62.5 mg/kg), leading to significant fibrotic lung disease within four weeks.

Genotyping was performed using primers listed in Supplementary Table S2, obtained from Invitrogen (Carlsbad, CA, USA) and IDT (Coralville, IA, USA). Mice were housed under specific pathogen-free conditions at the University of Cincinnati (UC), an American Association for the Accreditation of Laboratory Animal Care (AAALAC)-accredited facility. All animal experiments were conducted following protocols approved by the UC Institutional Animal Care and Use Committee (IACUC).

### Human Lung Sample Collection and Single Nuclei Isolation

Frozen lung tissues were collected from the basal segments of the right lower lobe and carefully excised to obtain approximately 50 mg of tissue free of visible airway structures, vessels, mucin, or blood clots. Tissues were immediately placed into a nuclei isolation buffer (20 mM Tris-HCl pH 7.5, 292 mM NaCl, 2 mM CaCl2, 42 mM MgCl2, 1% CHAPS, and 2% BSA) and homogenized using a Dounce homogenizer to mechanically dissociate nuclei from cells. The resulting homogenate was filtered through a 40 µm strainer, and nuclei were pelleted by centrifugation at 300 × g for 2 minutes at 4°C. Nuclei were resuspended in 0.04% BSA buffer to a final concentration of 1,000 nuclei/µl. Single-nuclei RNA sequencing (snRNA-seq) libraries were prepared using the 10x Genomics 5’ Kit v2, following the manufacturer’s protocol (10X Genomics). Libraries were sequenced on an Illumina NextSeq 550 platform using 75-cycle paired-end sequencing.

### Data Processing and Quality Control

Raw sequencing data were demultiplexed using Illumina’s bcl2fastq v2.20.0.422 software with default parameters, generating individual fastq files for each sample. These fastq files were aligned to the human reference genome (hg38) on 10x Genomics Cell Ranger v7.0.1 using 10x Genomics Cloud Analysis, which generated cell barcode, feature, and matrix files. Data were further processed in R Studio (v2023.03.0) using the Seurat package (v4.3.0) (46). Individual sample Seurat objects were created using “.h5” files, where Seurat objects went through quality control scrutiny before final analysis. Low-quality cells were first filtered below the threshold of 1000 detected genes and having greater than 20% mitochondrial gene expression per cell (22). Later, doublets were identified and removed using DoubletFinder (v2.0.3) (47), where artificial doublets are created based on preliminary clustering and cell clusters around the artificial clusters are classified as Doublets.

### snRNA-Seq Analysis

All analyses were performed using standard Seurat functions. The merged object was first log-normalized to transform the gene expression values, where raw counts were normalized and scaled up by a factor of 1000, and followed by log-transformed. Subsequently, 3000 variable genes were identified for each individual object using the Find Variable Features function. For these identified genes, data was scaled to have a zero mean across all cells and a variance of one using the Scale Data function. We used Run PCA function and identified thirty principal components (PCs) and used them to identify nearest neighbors using the Find Neighbors function, and clusters using the Find Clusters function. The resolution was adjusted to identify the predicted number of cell types. The reciprocal PCA protocol was adapted to integrate multiple Seurat objects and remove batch effects while preserving biological identity. UMAP plots were used to visualize clustering with a resolution of 0.7 for all cells and 0.3 for major cell types using the RunUMAP function. For data analysis and visualizations using heatmaps and violin plots, data in the RNA slot was used.

### Cell Type Annotation and Differential Gene Expression Analysis

Seurat-generated cell clusters were categorized into four major cell populations based on canonical cell markers: COL1A2 for mesenchymal cells, EPCAM for epithelial cells, PECAM1 for endothelial cells, and PTPRC for immune cells (Supplementary Figure 1). Each major type was further subdivided into subclusters following the same pipeline described above and was manually annotated using known cell type markers (Supplementary Table S1). Multiplets, identified by the co-expression of markers from different cell types, were removed, and the final dataset was created by mapping all annotated subgroups (Figure 1C). To calculate the average gene expression for each cell type per sample, the Average Expression function was used, grouping cells either by sample alone or by both sample and cell type. Differentially expressed genes (DEGs) between cell types were identified using Seurat’s Find All Markers function with default parameters, which compared each cluster against all others to generate a gene list for each cell type. To assess differential expression between IPF and control lung cells, the Find Markers function was applied with default parameters. The Wilcoxon rank-sum test was used to generate p-values for all DEG analysis. Gene enrichment analysis was conducted using ToppFun application of the ToppGene suite (48), and results were visualized using Cytoscape v3.10.3 (49).

### Mouse model of bleomycin-induced pulmonary fibrosis

Bleomycin was prepared by mixing sterile bleomycin sulfate powder (Teva Parenteral Medicines, Irvine, CA) with sterile normal saline. For the repetitive bleomycin injury model, three doses of bleomycin were administered intratracheally at three-week intervals at a dose of 1.6 units per kg body weight in a total volume of 50 μL of sterile saline, and mice were euthanized at the end of the ninth week. For a single-dose bleomycin injury model, bleomycin was administered intratracheally at a dose of 3 units per kg bodyweight in a total volume of 50 μL of sterile saline, and mice were euthanized at the end of the third week.

To analyze lung function, mice were injected with 100 µl of pentobarbital (65 mg/ml) intraperitoneally. A cannula was inserted into the trachea by puncturing it ventrally, and lung function parameters were measured using Flexi Vent (SCIREQ Inc., Montreal, Canada), following established protocols (50). For biochemical, transcriptomic and pathologic analysis of lung tissues, animals were sacrificed, and the lungs were collected for histology, RNA, protein, and hydroxyproline.

### Hydroxyproline measurement

Total lung hydroxyproline levels were quantified using a colorimetry-based method (Catalog MAK008, Sigma-Aldrich, St. Louis, MO, USA) as described previously (51). Briefly, lung tissue was hydrolyzed with HCl and neutralized with sodium hydroxide. The neutralized hydroxyproline was oxidized to pyrrole followed by a reaction with Ehrlich’s reagent in perchloric acid, which forms a bright-colored chromophore, and measurement at a wavelength of 558 nm.

### Histology and Immunohistochemistry

Histology and immunostaining were performed as described previously (52). In brief, the paraffin-embedded lungs were sectioned at a thickness of 5 µm and stained with Masson’s trichrome. For immunostainings, lung sections were deparaffinized, and antigen retrieval was performed with 10 mM citric acid (pH 6.0). The sections were then incubated with 5 % donkey serum for 1 hour, followed by overnight incubation with species-specific primary antibodies. Next, the sections were incubated with species-specific secondary peroxidase antibodies and stained with DAB (3,3’-Diaminobenzidine). Finally, the lung sections were mounted, and visualized using Keyence BZ series microscopy, and the area of staining was quantified with the BZ-X analyzer.

### Multiplex fluorescent in situ-hybridization

Five-micrometer-thick paraffin-embedded lung sections from control and IPF lung tissues were hybridized with WT1 (Catalog 415581-C1) and ACTA2 (Catalog 444771-C2) probes according to the manufacturer’s instructions. In brief, lung sections were deparaffinized, pretreated with hydrogen peroxide, and treated with RNAscope Protease Plus for 15 min at RT. Probes and RNAscope signal amplifiers were applied. Finally, DAPI-stained lung sections were mounted, visualized using Leica STELLARIS 8 confocal microscopy, and analyzed with Leica Microsystems LAS X image analysis software.

### Primary lung fibroblast cultures

Adult human lung primary fibroblasts from control tissue, patients with IPF, and mice were isolated using collagenase digestion, as described previously (4), and then cultured in DMEM (10% FBS) for human fibroblasts or IMDM media for mouse fibroblasts (5% FBS). The culture medium was supplemented with glutamine, penicillin, streptomycin, amphotericin, and gentamycin. Lung resident fibroblasts were isolated with negative selection using anti-CD45 microbeads and passing the cells through magnetic columns. Fibroblasts used in this study were from passages 1 to 3.

### Transfection and Transduction of fibroblasts

For stealth siRNA-mediated studies, IPF fibroblasts were transfected with stealth siRNA control (Catalog 12935300 Invitrogen) or WT1-specific siRNA (Catalog HSS111388, Invitrogen) for 72 hours using a lipofectamine 3000 transfection kit (Life Technologies, Carlsbad, California) as described previously (10). Human primary fibroblasts were treated with a control adenovirus, or WT1 overexpressing adenoviral particles as described previously (53). Cells were harvested after 48–72 hours for RNA or protein lysates, as mentioned previously (10). For Cre-mediated in vitro experiments, TGFα^OE^WT1^fl/fl^ mice were fed with Dox food for 8 weeks, and the lungs cultured for five days to isolate lung resident fibroblasts as described above. The purified lung resident fibroblasts were transduced with either Cre-expressing adenovirus (AdCre) or control Adeno (Ad-Control) viral particles to excise loxP sites that flanked the WT1 gene for 72 hours followed by TUNEL assay, immunoblotting, and RNA isolation.

### Quantitative RT-PCR analysis

Total RNA was extracted from lung tissues and primary cells using the RNeasy kit (Qiagen Sciences, Valences, CA), with concentration determined using a Nanodrop2000 spectrophotometer (Thermo Scientific) as previously described (53). Reverse transcription was conducted using SuperScript III (Thermo Fisher Scientific), followed by real-time PCR using SYBR select master mix (Bio-Rad) and the CFX384 Touch Real-time PCR instrument (Bio-Rad), with analysis performed using CFX maestro software version 4.0. Target gene transcripts from mouse samples were normalized to hypoxanthine-guanine phosphoribosyl transferase (Hprt), while human transcripts were normalized to human beta-actin. RT-PCR primer for mice and human used in this study (Invitrogen, Carlsbad, CA, USA, and IDT, Coralville, IA, USA) are provided in Supplementary Table S3 and S4, respectively.

### Immunoblotting

Immunoblot analysis was performed as described previously (54). Briefly, total lung tissue lysates and primary cell lysates were prepared using the cell lysis buffer (Cell Signaling Technology, 9803S) with protease inhibitors cocktail (Sigma), and protein concentration was estimated using a BCA protein estimation kit (Thermo Fisher Scientific, Waltham, MA). After SDS-PAGE separation, proteins were transferred to nitrocellulose membrane and blocked with 5% BSA (MilliporeSigma; A9647) for 2 h at room temperature and probed with specific primary antibodies in blocking buffer at 4°C overnight, followed by detection with HRP-linked secondary antibodies. The band intensities were quantified using Image Lab software 6.1 (Bio-Rad, USA), with target proteins normalized to internal control GAPDH. The primary and secondary antibodies used in this study are listed with their dilutions in Supplementary Table S5.

### Measurement of collagen in fibroblast conditioned media

IPF fibroblasts were transfected with either control siRNA or WT1-specific siRNA and cultured for 72 hours in DMEM supplemented with 0.5% FBS. Following 72 hours of knockdown, conditioned media were collected and processed as previously described (55). Briefly, media samples were passed through a 0.45 μm filter to remove debris and subsequently concentrated using Amicon Ultra Centrifugal Filters (Thermo Scientific, Rockford, IL, USA). The concentrated media were then mixed with 1× SDS loading buffer, denatured at 98°C for 5 minutes, and subjected to Western blot analysis to detect collagen.

### BrdU cell proliferation assay

Cell proliferation was evaluated using the BrdU kit (Cell Signaling Technology, Denver, CO), as described previously (3). Briefly, lung resident fibroblasts isolated from control and cWT1^OE^ mice were cultured for 48 hours under low-serum conditions (0.5% serum), with BrdU labeling solution added after 24 hours in culture. Following 24 hours of BrdU labeling, cells were fixed, and BrdU immunodetection was performed according to the manufacturer’s instructions.

### TUNEL assay

Human or mouse fibroblasts were treated with siRNA using Lipofectamine 3000 transfection kit or transduced with adenovirus for 48 hours as described previously (53), followed by treatment with species-specific anti-Fas antibody (250 ng/mL, 05-201, clone CH11, Millipore) for 24 hours. Stealth negative control siRNA (Catalog 12935300 Invitrogen) and human WT1 stealth siRNA (Catalog HSS111388, Invitrogen) were purchased from Invitrogen. The cells were then fixed with 4% paraformaldehyde, and nuclei were stained using DAPI. Assessment of DNA fragmentation in apoptotic cells was conducted using the terminal deoxynucleotidyl transferase dUTP nick-end labeling (TUNEL) method with an In Situ Cell Death Detection kit, TMR red (Roche Diagnostics), following the manufacturer’s instructions. Imaging was performed at an original magnification of 20X using a Nikon AIR-A1 confocal microscope, and quantification was carried out using MetaMorph imaging software (v6.2; Molecular Devices).

### Immunofluorescence and confocal imaging

Immunofluorescence and confocal imaging were performed as previously described (9). Briefly, mouse lung tissues form control and cWT1^OE^ mice were fixed in 1:10 diluted formalin, embedded in paraffin, and sectioned at 5 µm thickness. Sections were deparaffinized followed by citric acid (pH 6.0) antigen retrieval, blocked using 5 % normal donkey serum and incubated with primary antibodies overnight at 4° C. The following day, the samples were incubated with species-specific secondary donkey Alexa Fluor (488 and 594) antibodies, and nuclei were stained with DAPI. Images were captured using a Leica STELLARIS 8 confocal microscope and analyzed using Leica Microsystems LAS X image analysis software. Antibody details and dilutions are provided in Supplemental Table S5.

## Statistical analysis

All data were analyzed using Prism (version 10.2.3 for Windows, GraphPad Software, San Diego, CA, USA). For multiple comparisons, one-way ANOVA with Tukey’s test was performed. Statistical significance between two groups was determined using Student’s two-tailed t-test. Data were presented as mean ± SEM to indicate variability. P values less than 0.05 were considered statistically significant.

## Data availability

Raw and processed sequencing data are uploaded to GEO (Accession number: GSE279404).

## Author contributions

HHE, VS, and SKM devised the project, were involved in designing and executing the experiments, analyzed data, and wrote the manuscript; PKP performed immunostaining; CPV, AGJ, HM, DP, and MBJ performed bioinformatic analysis and edited the manuscript; SKH, FXM and NG provided human lung tissues and edited the manuscript; CE and AS shared the WT1 transgenic mice and edited the manuscript.

## Acknowledgements

This study was supported in part by NIH 1R01HL134801 (SKM) and 1R01HL157176 (SKM).

