## Supplementary material for "Wilms’ tumor 1 impairs apoptotic clearance of fibroblasts in distal fibrotic lung lesions": Supplemetal data

### Supplementary figure 1

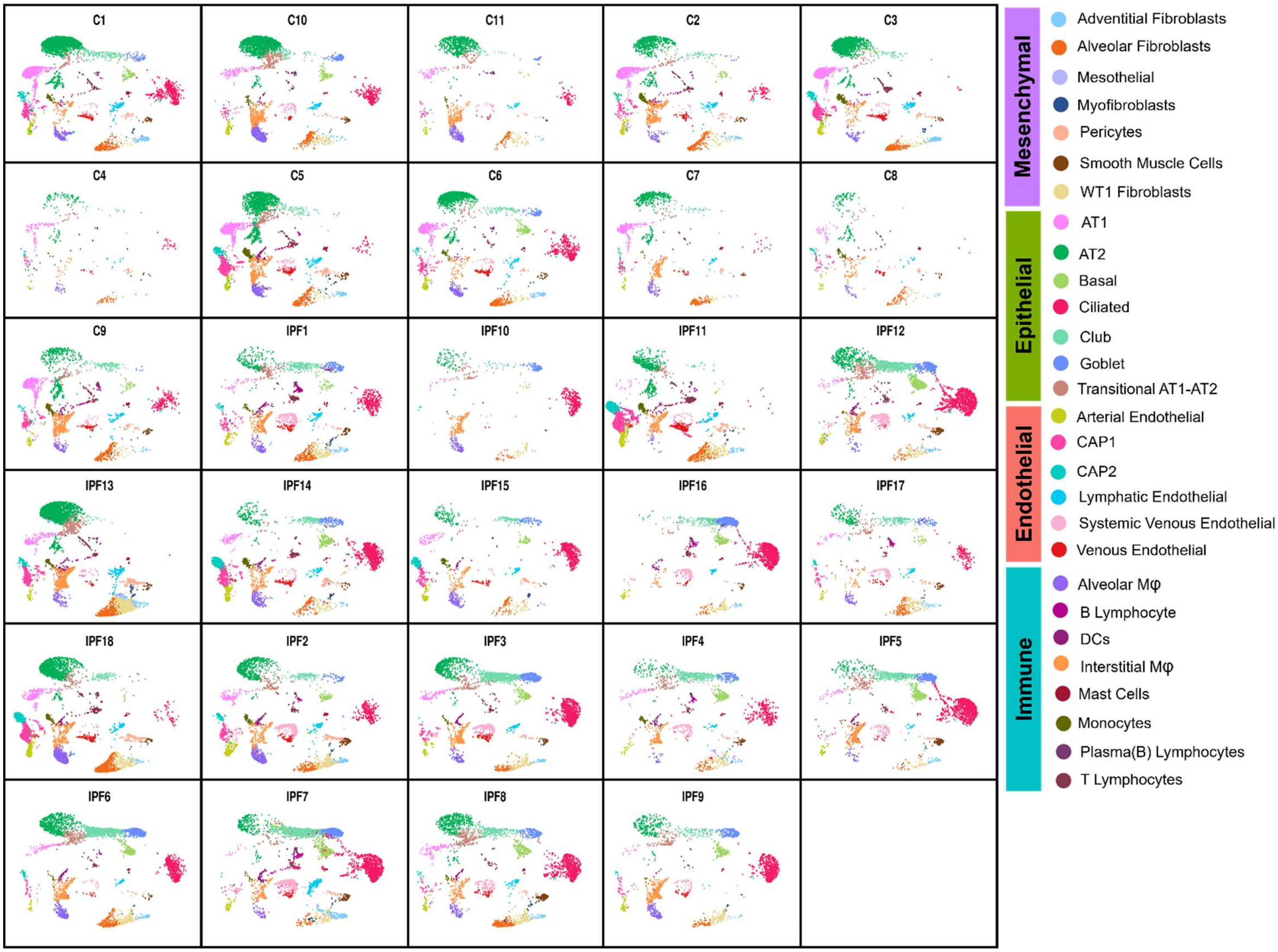


**Supplementary Figure 1.** The plots show UMAP layouts for all cell types from the control (N = 11, C1 to 11) and IPF (N = 18; IPF1 to IPF18) lung samples. Plots are colored by cell identity annotations.

### Supplementary figure 2

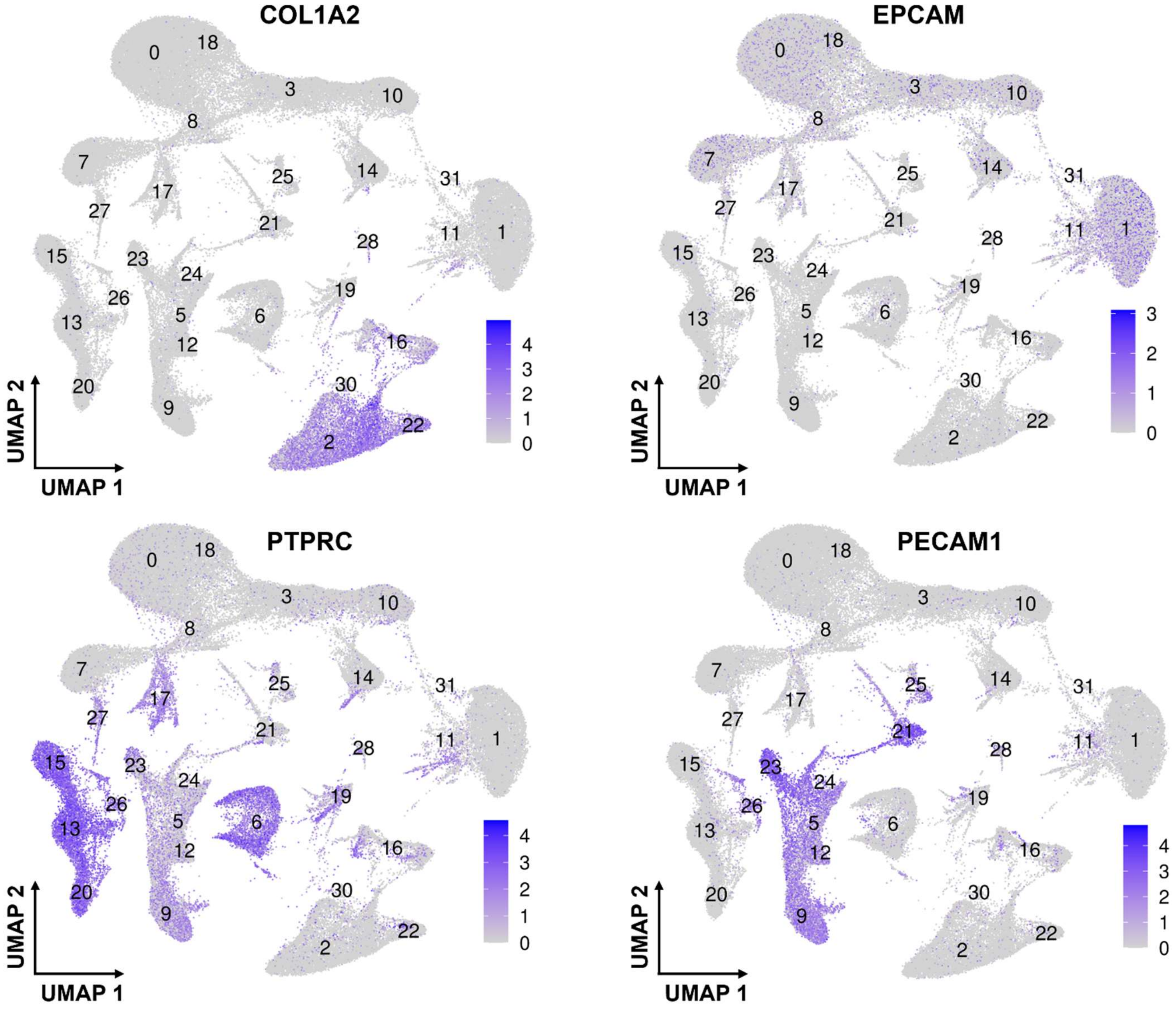


**Supplementary figure 2.** UMAP plots show the normalized gene expression of COL1A2, EPCAM, PTPRC, and PECAM1 of all cells for identifying mesenchymal, epithelial, immune, and endothelial cell populations, respectively. Color intensity reflects expression levels, with darker shades indicating higher expression.

### Supplementary figure 3

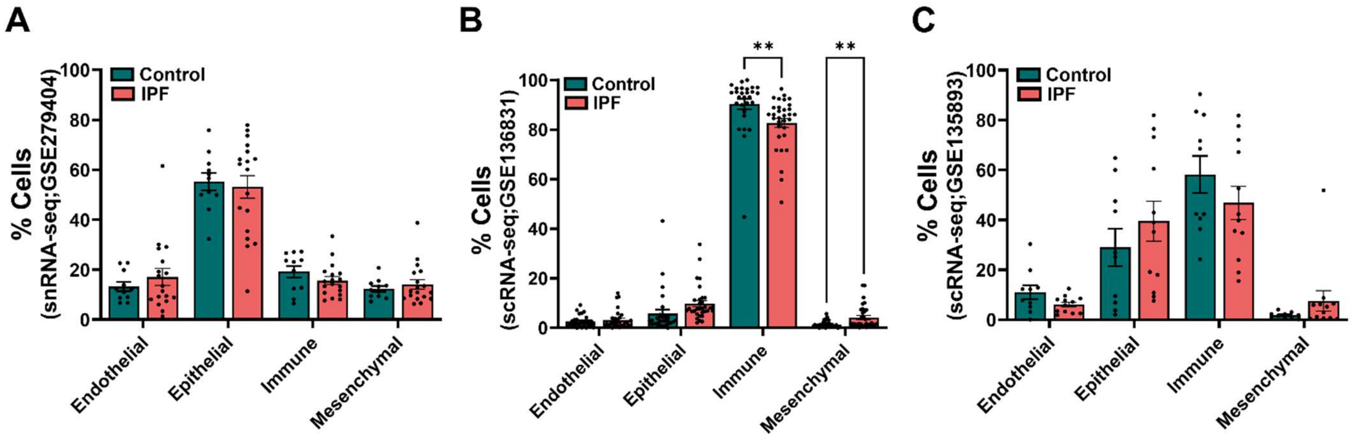


**Supplementary figure 3.** The percent distribution of all four major cell categories as a proportion of all sampled cells in control and IPF lungs across different single cell datasets, including **(A)** snRNA-seq (GSE279404), **(B)** scRNA-seq (GSE136831), and **(C)** scRNA-seq (GSE135893)**.** Each dot represents a single subject, and multiple t-test was used for comparisons between control and IPF (** $p < 0.01$).

**Supplementary figure 4**

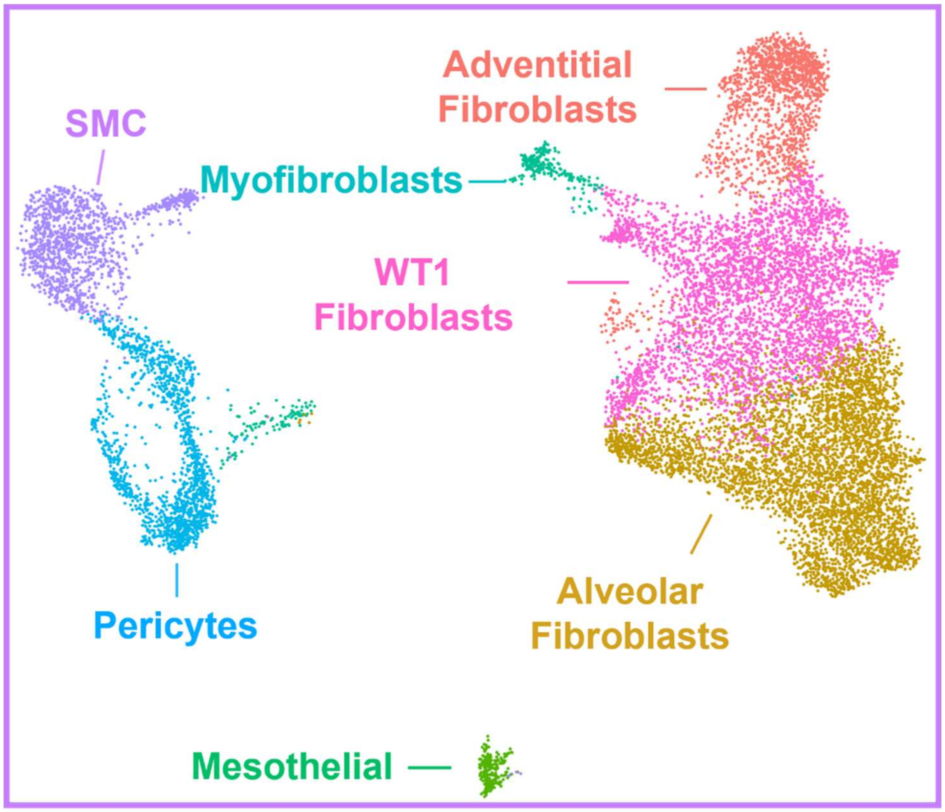


**Supplementary Figure 4.** UMAP plot shows color-coded 7 unique mesenchymal cell populations identified after the sub-clustering of all mesenchymal cells.

**Supplementary figure 5**

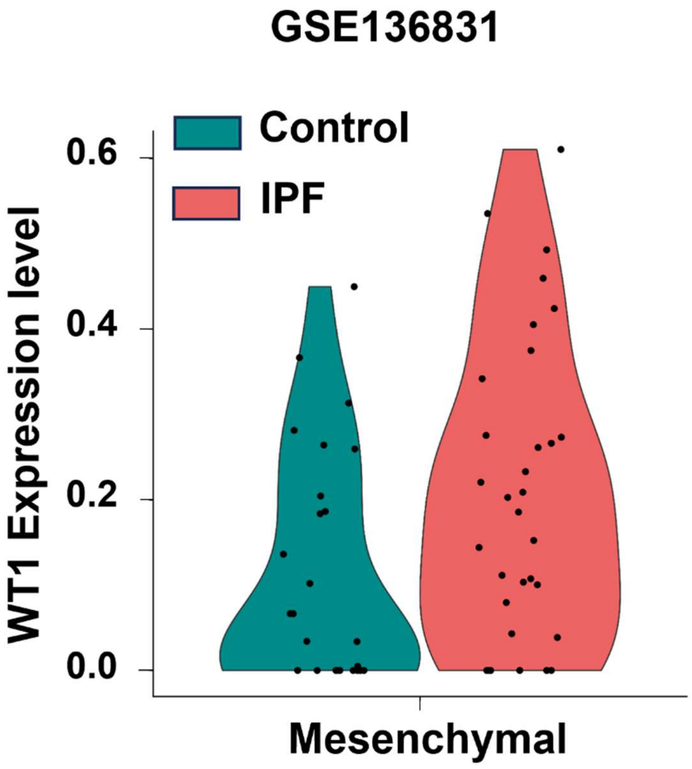


**Supplementary Figure 5.** Violin plot showing the average WT1 expression in the mesenchymal population comparing control and IPF subjects in a publicly available dataset (GSE136831). Each dot represents an individual subject. The p-value was calculated using the Wilcoxon rank-sum test (< 0.05).

### Supplementary figure 6

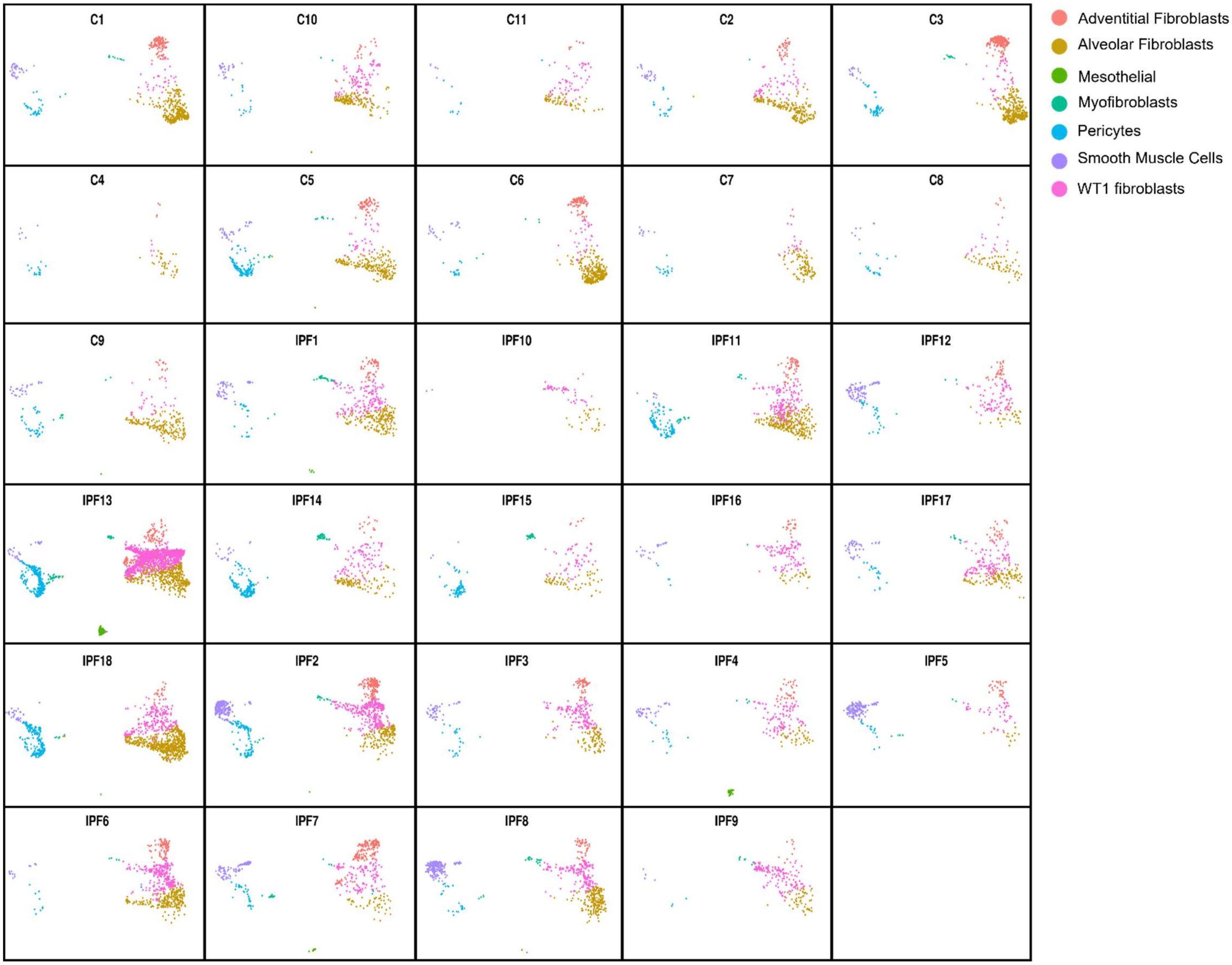


**Supplementary Figure 6.** The plots show UMAP layouts for mesenchymal cell populations from the control (N = 11, C1 to 11) and IPF (N = 18; IPF1 to IPF18) lung samples. Plots are colored by cell identity annotations.

**Supplementary figure 7**

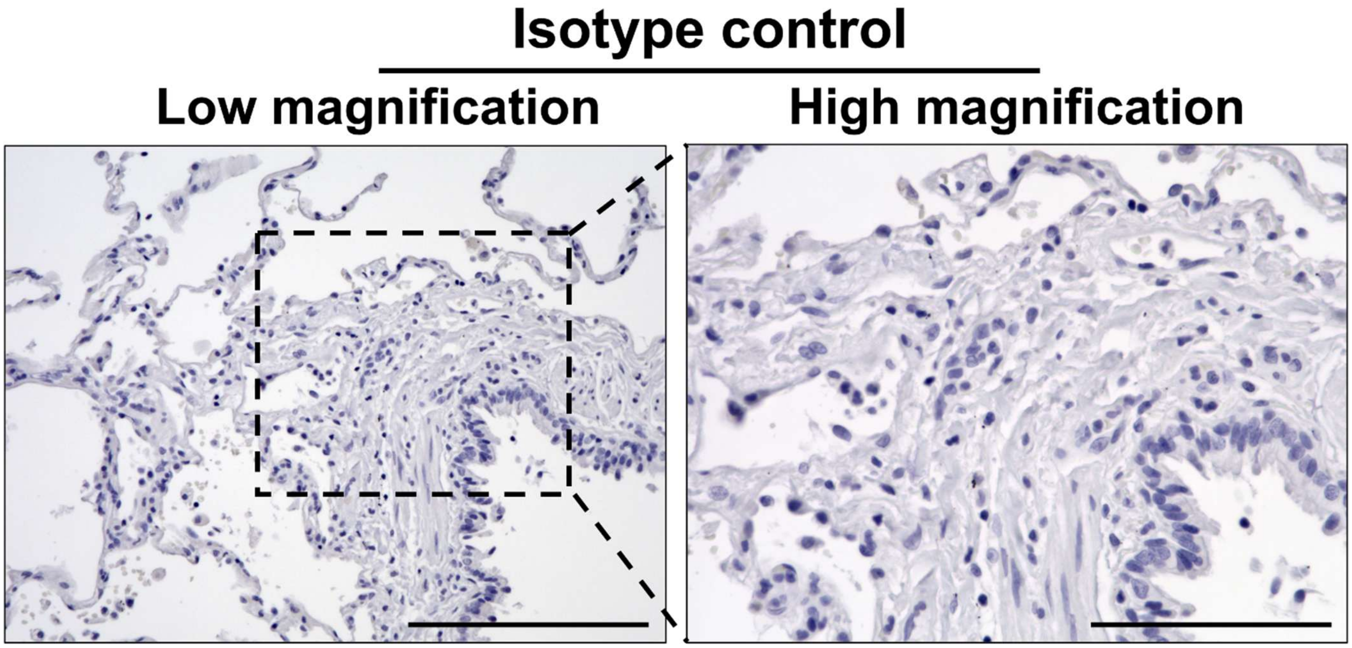


**Supplementary Figure 7.** Representative images of lung sections immunostained with an isotype control antibody (Rabbit IgG). Images were captured at low (20X; Scale bar, 200 µm) and high (40X; Scale bar, 100 µm).

**Supplementary figure 8**

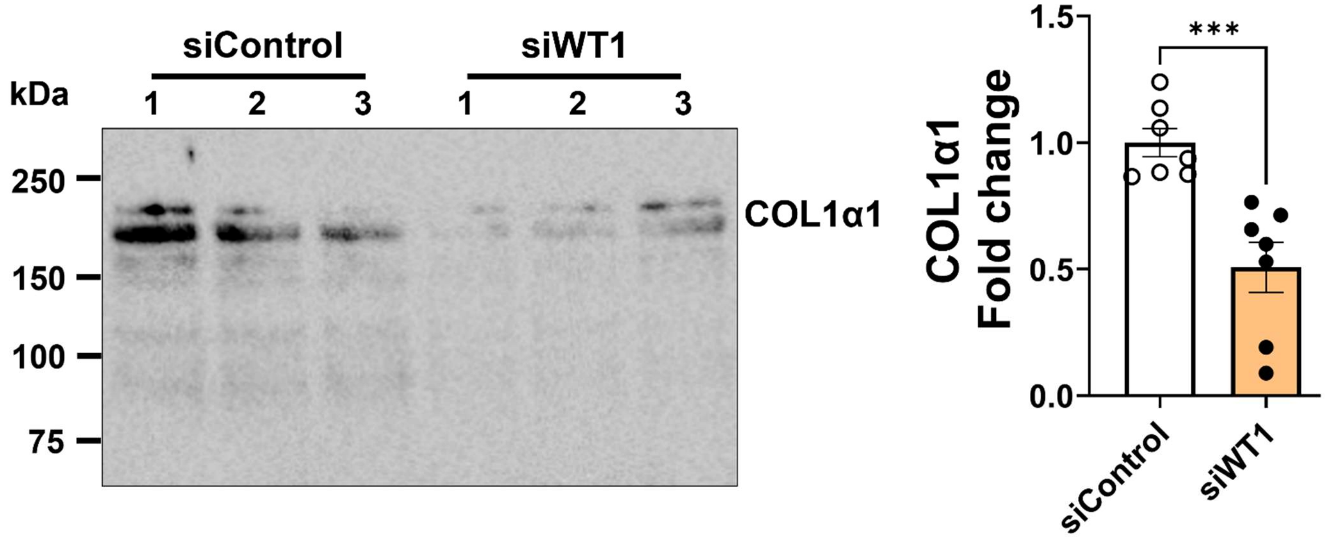


**Supplementary Figure 8. The loss of WT1 attenuates collagen secretion in IPF fibroblasts.** Conditioned media were collected from IPF fibroblasts treated with either control or WT1-specific siRNA for 72 hr and immunoblotted with an antibody against COL1α1. COL1α1 protein levels in conditioned media were shown as fold change normalized to total cells using a bar graph. Student's two-tailed *t*-test was used (** $p < 0.001$; n=7/group). Data is the cumulative result of two independent experiments with similar findings.

**Supplementary figure 9**

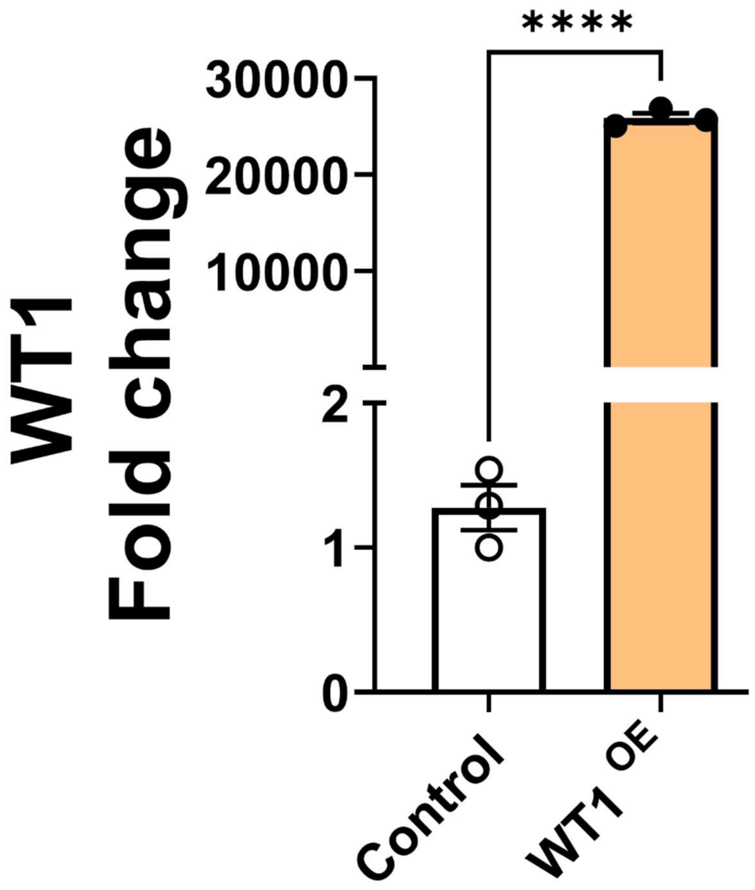


**Supplementary figure 9. Overexpression of WT1 using Adenovirus in normal lung fibroblasts.** Quantification of WT1 transcripts using RT-PCR in normal lung resident fibroblasts transduced with either control or WT1-Adeno viral particles for 72 hr. Student's two-tailed *t*-test was used (**** $p < 0.0001$; n=3/group).

Supplementary figure 10

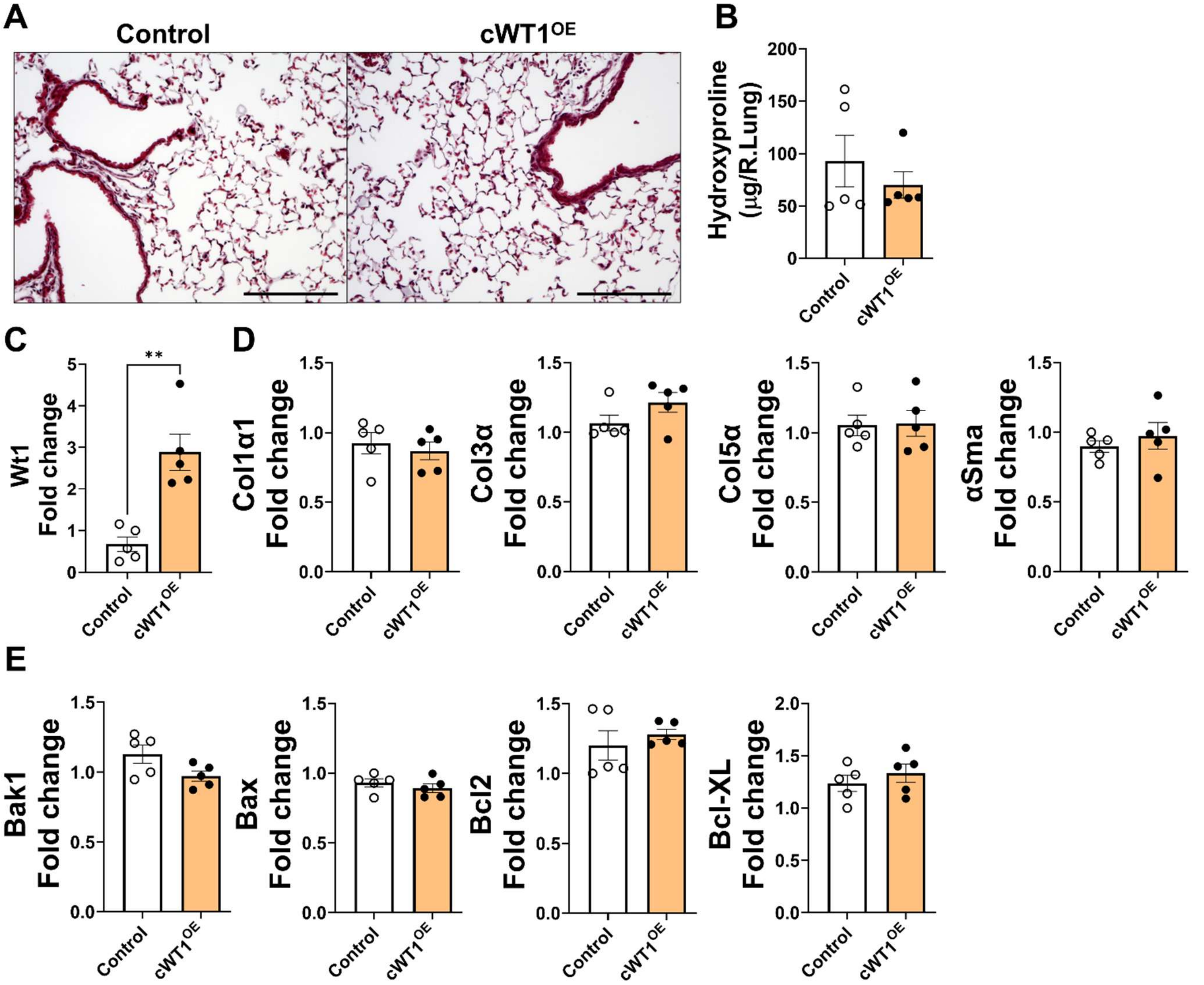


**Supplementary Figure 10.** (A) Masson's trichrome stained lung sections from young age control and cWT1$^{OE}$ mice treated with tamoxifen via intraperitoneal injection for two weeks. Images were captured at 20X original magnification. Scale bar, 200 µm. (B) Hydroxyproline levels were measured in the right lungs of control and cWT1$^{OE}$ mice treated with tamoxifen via intraperitoneal injection for two weeks (n=5/group). (C) Quantification of WT1 gene transcripts by RT-PCR in the total lung RNA isolated from control and cWT1$^{OE}$ mice treated with tamoxifen via intraperitoneal injection for two weeks. Student's two-tailed *t*-test was used (**$p < 0.01$;

n=5/group). (D-E) Quantification of ECM (Col1α1, Col3α, and Col5α) and apoptosis-associated gene transcripts (Bak1, Bax, Bcl2, and Bcl-XL) from control and cWT1$^{OE}$ mice treated with tamoxifen via intraperitoneal injection for two weeks.

**Supplementary figure 11**

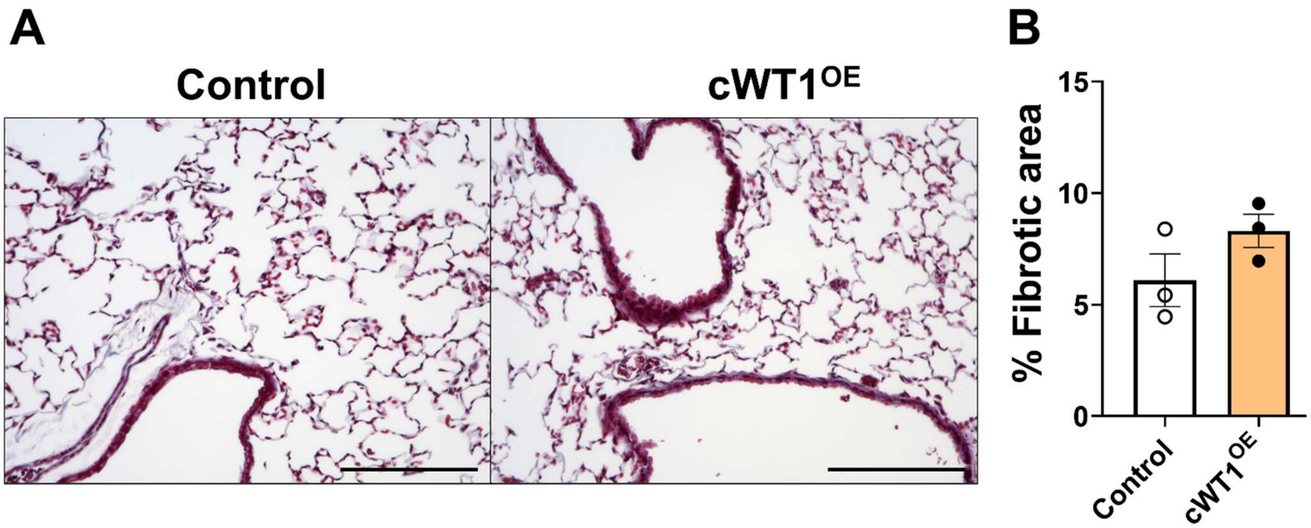


**Supplementary Figure 11.** (A) Masson's trichrome stained lung sections from old age control and cWT1$^{OE}$ mice treated with tamoxifen via intraperitoneal injection for two weeks. Images were captured at 20X original magnification. Scale bar, 200 μm. (B) Percent fibrotic area was quantified in control and cWT1$^{OE}$ mice using BZ-X image analysis. (n=3/group).

**Supplementary figure 12**

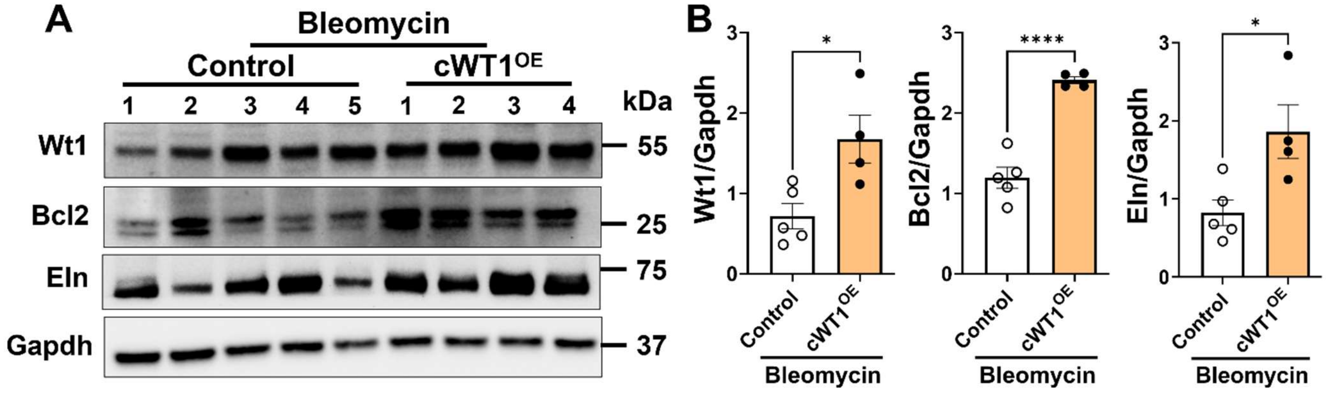


**Supplementary Figure 12. WT1 increases the molecular markers of survival and ECM during bleomycin-induced pulmonary fibrosis.** The total lung lysates were immunoblotted with antibodies against Wt1, Bcl2, Eln, and Gapdh from old age control and cWT1$^{OE}$ mice treated with bleomycin. Wt1, Bcl2, and Eln protein levels were normalized to Gapdh and shown as fold change using a bar graph. Student's two-tailed *t*-test was used (* $p < 0.05$; **** $p < 0.0001$; n=4-5/group).

**Supplementary Table S1.** Markers used for cell annotation in snRNA-seq analysis.

| Population | Cell Type Annotation | Marker(s) |
|---|---|---|
| Mesenchymal | Alveolar Fibroblasts | MYLK, TCF21, ADGRB3, DCC, LIMCH1 |
| Mesenchymal | Adventitial Fibroblasts | MFAP5, SCARA5, IGF1, SPOCK1 |
| Mesenchymal | Myofibroblasts | DACH2, ANO4, ITGBL1 |
| Mesenchymal | WT1 fibroblasts | WT1, LUM, COL1A1, POSTN, COL3A1 |
| Mesenchymal | Mesothelial | WT1, MSLN, CA12, MIR31HG, RBFOX1 |
| Mesenchymal | SMCs | MYH11, ACTA2, LGR6, LDB3 |
| Mesenchymal | Pericytes | LAMC3, TRPC6, PDGFRB, DGKG |
| Endothelial | Arterial Endothelial | GJA5, DKK2, ENPP2, ARL15 |
| Endothelial | Lymphatic Endothelial | PROX1, MMRN1, CCL21, RELN, DLGAP2 |
| Endothelial | Venous Endothelial | ACKR1, SULT1E1, HDAC9, ADGRG6, FAM155A |
| Endothelial | Systemic Venous Endothelial | COL15A1, VWA1, ESR2, MYRIP, POSTN |
| Endothelial | CAP1 | APLNR, GPIHBP1, IL7R, FCN3, NRXN3 |
| Endothelial | CAP2 | CAR4, APLN, EDNRB, HPGD, TBX2 |
| Epithelium | AT1 | AGER, RTKN2, NCKAP5, EMP2, SCEL |
| Epithelium | AT2 | SFTPC, LAMP3, LRRK2, ABCA3, SFTPB |
| Epithelium | Basal | KRT5, TP63, MIR205HG, KRT15, EYA2 |
| Epithelium | Ciliated | FOXJ1, RSPH1, PTPRT, DCDC1, DNAH12 |
| Epithelium | Club | SCGB3A2, HMCN1, SLC4A5 |
| Epithelium | Goblet | MUC5AC, SPDEF, MUC5B, SCGB1A1, SCGB3A1 |
| Epithelium | Translational AT1-AT2 | Express both AT1 (AGER, RTKN2, NCKAP5, EMP2, SCEL) and AT2 markers (SFTPC, LAMP3, LRRK2, ABCA3, SFTPB) |
| Immune | Alveolar Macrophages | MSR1, MCEMP1, PPARG, LSAMP |
| Immune | Interstitial Macrophages | F13A1, STAB1, FMN1, ABCA1 |
| Immune | Monocytes | VCAN, S100A8, LILRB2, LILRA5 |
| Immune | B Lymphocytes | MS4A1, CD19, BLK, BACH2 |
| Immune | Plasma (B) Lymphocytes | FCRL5, JCHAIN, TXNDC5, IGKC |
| Immune | T Lymphocytes | CD3E, IL7R, ITK, CD247 |
| Immune | DCs | HDAC9, DC1, FCGR2B, CLEC10A, PALD1 |
| Immune | Mast Cells | CPA3, TPSD1, GATA2, MS4A2 |

**Supplementary Table S2.** The list of mouse genotyping primers used in the study.

| Gene Symbol | Forward primer | Reverse primer |
|---|---|---|
| CC10 | ACTGCCCATTGCCCAAACAC | AAAATCTTGCCAGCTTTCCCC |
| TetO | CCTGTTCGCTCTGGGTATTGTGTT | CGTGGTCCGCTGATTTCTTCTCTA |
| PDGFRα CreERT | TCA GCC TTA AGC TGG GAC AT | ATG TTT AGC TGG CCC AAA TG |
| $WT1^{fl/fl}$ | GCC CTA CAG CAG GTA AGA AGG | CTG GAG ACC TGA GAC AAG CA |
| $WT1^{OE}$ | CACCAAAGGAGACACACAGGT | GGGCTTTTCACCTGTATGAG |

**Supplementary Table S3.** The list of mouse RT-PCR primers used in the study.

| Gene Symbol | Forward primer | Reverse primer |
|---|---|---|
| Acta2 | TGACGCTGAAGTATCCGATAGA | CGAAGCTCGTTATAGAAAGAGTGG |
| Bad | GGAGCAACATTCATCAGCAG | TACGAACTGTGGCGACTCC |
| Bak1 | GGAATGCCTACGAACTCTTCA | CCAGCTGATGCCACTCTTAAA |
| Col1α1 | CATGTTCAGCTTTGTGGACCT | GCAGCTGACTTCAGGGATGT |
| Col1α2 | AAGGGTGCTACTGGACTCCC | TTGTTACCGGATTCTCCTTTGG |
| Col3α1 | TCCCCTGGAATCTGTGAATC | TGAGTCGAATTGGGGAGAAT |
| Col5α | CTACATCCGTGCCCTGGT | CCAGCACCGTCTTCTGGTAG |
| Eln | TGGAGCAGGACTTGGAGGT | CCTCCAGCACCATACTTAGCA |
| Fn1 | CGGAGAGAGTGCCCCTACTA | CGATATTGGTGAATCGCAGA |
| Hprt | GCCCTTGACTATAATGAGTACTTCAGG | TTCAACTTGCGCTCATCTTAGG |
| Plk1 | TTGTAGTTTTGGAGCTCTGTCG | CAGTGCCTTCCTCCTCTTGT |
| Wt1 | CAG ATG AAC CTA GGA GCT ACC TTA AA | TGC CCT TCT GTC CAT TTC A |

**Supplementary Table S4.** The list of human RT-PCR primers used in the study.

| Gene Symbol | Forward primer | Reverse primer |
|---|---|---|
| ACTA2 | GCTTTCAGCTTCCCTGAACA | GGAGCTGCTTCACAGGATTC |
| β-ACTIN | CCAACCGCGAGAAGATGA | CCAGAGGCGTACAGGGATAG |
| BAX | AGCAAACTGGTGCTCAAGG | CTTGGATCCAGCCCAACA |
| BIM | ACTGGAGAGCTCATTGCAGAC | AAATACCAGGACCCGAAGGT |
| BCL2 | AGTACCTGAACCGGCACCT | GCCGTACAGTTCCACAAAGG |
| BCL2-L2 | TGGATGGTGGCCTACCTG | CGTCCCCGTATAGAGCTGTG |
| BCL-XL | GCCACTTACCTGAATGACCAC | TGCTGCATTGTTCCCATAGA |
| BCL3 | GCCTCAGCTCCAATGGTC | GAGGAGCCATGGGGAATC |
| COL1α1 | GGGATTCCCTGGACCTAAAG | GGAACACCTCGCTCTCCA |
| COL6α3 | ATGAGGAAACATCGGCACTTG | GGGCATGAGTTGTAGGAAAGC |
| COL16α1 | CCACCAGAAGACGTGGTATCT | CAGGACACAAAGTCGCCATC |
| HSP90B1 | GCTGACGATGAAGTTGATGTGG | CATCCGTCCTTGATCCTTCTCTA |
| PIM1 | GGCTCGGTCTACTCAGGCA | GGAAATCCGGTCCTTCTCCAC |
| WT1 | AGCTGTCCCACTTACAGATGC | CCTTGAAGTCACACTGGTATGG |

**Supplementary Table S5.** The list of antibodies and their dilutions used in the study.

| Antibody | Dilution | | | Catalog # | Company |
|---|---|---|---|---|---|
| | IHC | IF | WB | | |
| ACTA2 | 1:20000 | 1:2000 | 1:20000 | A5228 | Sigma |
| BCL-XL | | | 1:1000 | 2764S | CST |
| BCL2 | | | 1:1000 | 3498S | CST |
| BAD | | | 1:1000 | 9292S | CST |
| BAK | | | 1:1000 | 12105S | CST |
| BAX | | | 1:1000 | 2772S | CST |
| COL1α1 | | | 1:1000 | 91144S | CST |
| COL1α1 | | | 1:500 | Sc-293182 | Santa Cruz Biotechnology |
| ELN | | | 1:500 | 15257-1-AP | Proteintech |
| FN1 | | | 1:500 | Sc-9068 | Santa Cruz Biotechnology |
| FAS | | | 1:1000 | 8023S | CST |
| GAPDH | | | 1:2000 | A300-641 | Bethyl Laboratories |
| Vimentin | | 1:50 | | Sc-7557 | Santa Cruz Biotechnology |
| WT1 | 1:200 | 1:50 | | Ab89901 | Abcam |
| WT1 | | | 1:500 | 12609-1-AP | Proteintech |
